# Lentiviral single-cell MPRA of synthetic enhancers reveals motif affinity-based encoding of cell type specificity

**DOI:** 10.64898/2026.02.27.708495

**Authors:** Julia Rühle, Robert Frömel, Aina Bernal Martinez, Chelsea Szu-Tu, Joseph Bowness, Lars Velten

**Affiliations:** Computational Biology and Health Genomics, Centre for Genomic Regulation (CRG), Barcelona, Spain; Universitat Pompeu Fabra (UPF), Barcelona, Spain; Present address: Research Unit Brain Epigenomics, Helmholtz Center Munich, 81377 Munich, Germany

## Abstract

Cell-state-specific gene expression programs emerge from the interplay between cis-regulatory elements (CREs), such as enhancers, and transcription factors (TFs). Massively parallel reporter assays (MPRAs) have enabled large-scale dissection of CRE design principles, but bulk approaches cannot resolve cell state-specific regulatory logic on continuous trajectories of cellular differentiation, and existing single-cell MPRAs are not readily applicable to primary cell differentiation models. Here, we developed a single-cell lentiviral Massively Parallel Reporter Assay (sc-lentiMPRA) that overcomes these limitations and enables parallel quantification of enhancer activity and cellular transcriptome. Applying sc-lentiMPRA in blood stem differentiation, we profiled the activity and specificity of ~160 fully synthetic enhancers with controlled motif composition and affinities across ~190,000 single cells. Focusing on Trp53 and Cebpa, we show that enhancers with high and low affinity motifs differ qualitatively and quantitatively in their responses to TF expression gradients. For Trp53, low-affinity motifs exhibited near-linear correlation with TF expression, whereas high-affinity motifs showed reduced sensitivity to TF levels and a potential contribution of cofactor availability. In contrast, Cebpa-associated enhancers displayed nonlinear behaviors. Together, sc-lentiMPRA establishes a powerful framework for systematically relating enhancer architecture and TF expression to regulatory output at single-cell resolution during cellular differentiation.

## Background

During cellular differentiation, gradual and often overlapping patterns of transcription factor (TF) expression are converted into precise on/off activities of cell identity genes and transcriptional programs. In the example of blood stem cell differentiation (hematopoiesis), single-cell RNA-seq [1–3] and imaging-based [4] studies have demonstrated that master transcriptional regulators are expressed throughout the early stem- and progenitor compartment, whereas the expression patterns of downstream effectors are much more specific. These observations, together with related findings in other systems, raise the fundamental question of how TF expression levels and cis-regulatory elements (CREs), such as enhancers, interact to achieve cell state specificity.

Massively Parallel Reporter Assays (MPRAs) enable the functional dissection of CREs by systematic, high-throughput activity measurements of hundreds of thousands of natural or synthetic DNA sequences in parallel [5–8]. In MPRA, candidate CREs are cloned upstream or downstream of a reporter gene with a minimal promoter, and CRE activity is quantified by next-generation sequencing. Applications of MPRAs include studies of disease-associated variants [9–11], enhancer engineering for gene therapy [12], as well as work on general principles of gene regulation [13–15]. However, many existing MPRA datasets were acquired in immortalized cell lines [5–8] or flow-sorted, yet heterogeneous primary cell populations [10,16]. They are therefore limited in their ability to investigate how cell state-specific enhancer activity is achieved, and more specifically, to relate enhancer activity with TF expression levels during cellular differentiation. Recently, there has been an interest in developing single cell variants of MPRAs that can overcome this limitation [17–20]. On a technical level, existing single-cell assays are based on transfection and therefore limited in their compatibility with primary cell studies and many cellular differentiation models, including hematopoiesis. Furthermore, existing assays are based on whole transcriptome sequencing, which compromises capture rates of transcripts used for enhancer quantification, and increases cost.

While most MPRAs use variants of natural regulatory sequences, MPRAs of fully synthetic, minimalist enhancers have emerged as a powerful tool to characterize enhancer design principles. In synthetic MPRAs, the effect of single TF binding sites (motifs), or combinations of motifs, on transcriptional regulation can be studied outside of native, and often complex, DNA context. For example, we have recently employed a design where candidate motifs were placed in different random DNA background sequences [16], enabling precise control over motif number, motif affinity, orientation, and spacing between motifs. In hematopoietic stem and progenitor cells (HSPCs), we thereby discovered TF motifs that changed their function from activators to repressors of transcription as a function of site number (“occupancy-dependent duality”) or when combined with sites for other TFs (“combinatorial duality”). Finally, the use of synthetic sequences in MPRA is also supported by the theoretical argument that the genome is simply too small and redundant to capture the full spectrum of regulatory rules and the combinatorial complexity of TF interactions [21]. Characterizing these fully synthetic enhancers in cellular differentiation at the single-cell level has the potential to systematically link enhancer design features, TF expression levels, and enhancer activity at high resolution.

Here, we establish a lentiviral single-cell Massively Parallel Reporter Assay (sc-lentiMPRA) to dissect gene regulatory logic along developmental trajectories. We first addressed limitations of existing single-cell MPRA by developing a lentiviral implementation that enables reproducible high-throughput profiling of enhancer activity and a cost-effective, targeted transcriptome readout [22]. We applied this method to characterize the activity and specificity of minimal synthetic enhancers [16] in *ex vivo* HSPC differentiation and stem cell expansion cultures [16,23]. Specifically, we designed enhancers containing random background DNA and varying numbers of TF binding sites, at varying affinity, with a focus on the TFs Trp53 and Cebpa. For Trp53, we found that low-affinity binding sites linearly react to changes in TF expression, whereas high-affinity motifs show reduced dosage dependence, and appeared regulated by cofactor availability. In contrast, Cebpa-associated enhancers exhibited more complex behavior, indicative of non-linear interactions. These examples illustrate the potential of sc-lentiMPRA.

## Results

### Design of a lentiviral single-cell MPRA

We developed a lentiviral single-cell Massively Parallel Reporter Assay (sc-lentiMPRA) framework to profile synthetic enhancer activity in individual cells across the HSPC differentiation landscape. sc-lentiMPRA is based on the lentiMPRA vector [24,25], which uses anti-repressor elements (ARE) to minimize integration site effects. Our sc-lentiMPRA method enables joint measurement of reporter activity and gene expression at the single-cell level, offering a high-throughput approach to investigate enhancer function in dynamic and heterogeneous primary cell contexts.

In bulk lentiMPRA, enhancer activity is quantified by sequencing barcoded reporter mRNAs and normalizing their counts to the corresponding DNA barcode counts, which measure differences in CRE representation due to library composition and delivery. At the single-cell level, combined DNA/RNA readouts are possible [26–28], but low in throughput or impractical in an MPRA setting. Importantly, reporter mRNA alone is insufficient to interpret low or absent activity in a single-cell setting, because it cannot distinguish between a CRE that is not present in a cell and one that is present but inactive; thus, measuring construct presence (i.e., CRE representation) in each cell is essential. We therefore designed a vector that encodes both enhancer presence and activity within RNA transcripts [17,18]. Our sc-lentiMPRA vector features two independent transcriptional units. A “presence barcode” is constitutively expressed from a Pol III-dependent hU6 promoter via a gRNA scaffold [29], which serves solely as the carrier of this barcode and has no targeting role. A second “quantification barcode” is embedded in the 5′ UTR of GFP, downstream of the putative enhancer and a minimal Pol II-dependent promoter (Fig. 1a). The use of distinct Pol II and Pol III promoters minimizes cross-talk between the two expression cassettes. Specifically, orienting the hu6 and minimal promoter expression cassettes in the same transcriptional direction (head-to-tail), showed higher independence compared to a head-to-head orientation (Additional File 1: Fig. S1). Additionally, our design minimized barcode swapping by lentiviral recombination through close physical proximity between the two barcodes (Additional File 1: Fig. S2).

**Fig. 1:**
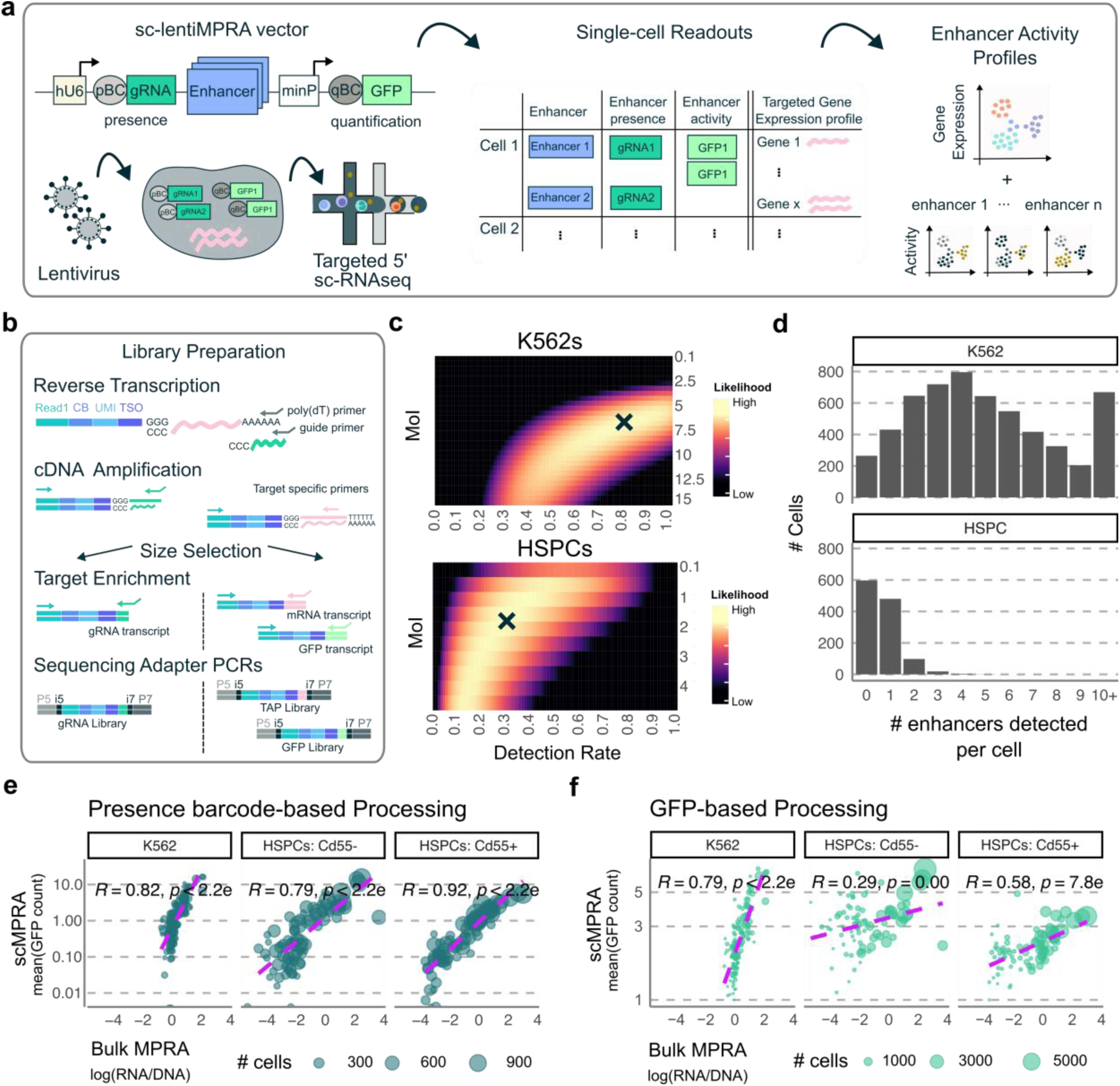
sc-lentiMPRA enables single-cell measurement of enhancer activity. **a** Schematic of the sc-lentiMPRA workflow combining lentiviral delivery of an enhancer library with 5′ single-cell RNA-seq to jointly profile enhancer activity and targeted transcriptomes. **b** Library preparation strategy generating separate sequencing libraries for presence barcodes (gRNAs), quantification barcodes (GFP), and targeted transcriptome (TAP) [22]. **c** Estimated probability of detecting enhancer-presence barcodes per cell by fitting a model [32,33] to the distribution of observed guide barcodes as function of multiplicity of infection (MoI) and barcode capture probability in K562 cells and primary HSPCs. x indicates the maximum likelihood. Distribution of detected enhancers per cell in K562 cells and HSPCs. **e** Correlation between bulk MPRA and mean single-cell enhancer activities in K562 cells and two HSPC populations (Cd55^+^ and Cd55^−^) using sc-lentiMPRA preprocessing that uses presence and quantification barcodes (Pearson’s R = 0.79–0.92). **f** Correlation between bulk MPRA and mean single-cell enhancer activities in K562 cells and two HSPC populations (Cd55^+^ erythromyeloid and Cd55^−^ monocyte-neutrophil progenitors [16]) only using the GFP-based quantification barcode for enhancer activity quantification (Pearson’s R = 0.29–0.79).

To translate this vector design into a robust and scalable sc-lentiMPRA workflow, we optimized a cloning protocol that achieves even coverage of enhancer libraries and adjustable barcode complexities, and developed a sequencing strategy to associate barcodes and CREs (Additional File 1: Fig. S3, S4). In sum, we designed a scMPRA vector for lentiviral delivery that includes two independently expressed barcodes for sensitive quantification of enhancer activity at the single - cell level. Our design decreases lentiviral recombination rate while avoiding transcriptional interference between detection and quantification reporters.

### Validation of sc-lentiMPRA

To validate the sc-lentiMPRA framework, we designed a Proof-of-Concept (PoC) library of 432 synthetic and natural genomic elements that covered a wide range of activities in bulk MPRA experiments ([16] and see Methods), and performed a pilot experiment in both K562 cells and differentiation cultures derived from primary mouse Hematopoietic Stem Cells (HSCs). For single-cell readouts, we developed a custom 5’ scRNA-seq protocol (modified from the Chromium Next GEM Single Cell 5’ kit from 10x Genomics) that enables a) direct capture of the gRNA transcript that carries the presence barcode [30], b) PCR-based targeted capture of a panel of mRNAs [22], and c) enrichment for the GFP reporter transcript (Fig. 1b and Methods for protocol detail, Additional File 2). For the K562 experiment, we used a simple targeted panel of 30 genes, while for the HSPC differentiation culture, we employed an expanded panel consisting of 287 primers tailored to cover key TFs and lineage-defining transcripts (Methods, Additional File 3). We found that expression of a nuclease-dead dCas9 construct in the cells of interest was required to stabilize the gRNA transcript and permit efficient capture.

After preprocessing and cell quality control (see Methods), we evaluated the performance of the sc-lentiMPRA method at the level of transcriptome, guide and GFP barcode readouts. For the targeted transcriptome capture, we observed a clear enrichment of targeted over non-targeted genes, with high UMI counts per cell (median = 398 in K562s and median = 1,198 in HSPCs), despite using < 2000 reads per cell (Additional File 1: Fig. S5). This demonstrates the effectiveness of the targeted capture approach. At the level of the gRNAs, we first established thresholds to minimize false positive enhancer calls ([31] and see Methods). We then estimated the capture rate for the presence barcodes by modeling the distribution of observed guide barcodes per cell as a function of multiplicity of infection (MoI) and capture probability [32,33]. This analysis revealed high capture efficiencies of ~80% in K562 cells and ~30% in HSPCs, at estimated MoIs of 6.6 and 1.6, respectively (Fig. 1c). This resulted in 30,194 enhancer observations in 5,666 K562 cells with ~95 % of the cells containing at least one enhancer (presence barcode), and 754 enhancer observations in 1199 profiled HSPCs, with enhancer detection in ~50 % of the cells (Fig. 1d).

To quantify enhancer activity, we first assigned each cell to enhancer construct(s) based on detection of presence barcodes. We then quantified activity of the respective enhancer as the number of GFP (quantification barcode) UMI counts (see Methods). GFP UMI count per cell, gRNA UMI count per cell and total targeted transcriptome mRNA UMI count per cell showed negligible correlations of Library PoC enhancers in K562 cells (R<0.05), indicating that technical effects or differences in mRNA content between cells do not bias enhancer activity estimates (Additional File 1: Fig. S6). To validate the sc-lentiMPRA-derived enhancer activity estimates we compared sc-lentiMPRA “pseudobulks” with bulk lentiMPRA [24] measurements from K562 cells and from primary HSPCs flow-sorted into a CD55^+^ (GATA2-positive erythromyeloid progenitors) and a CD55^−^ (GATA2-negative monocyte and neutrophil progenitors) population, and observed strong correlations (R = 0.79 - 0.92) (Fig. 1e, Additional File 1: S7a,b). For comparison, we evaluated a preprocessing strategy that omitted the presence barcode and relied solely on the quantification GFP barcode, resulting in lower correlations (R = 0.29-0.79) (Fig. 1f). This analysis showed that use of the presence barcode consistently improved agreement with bulk MPRA measurements. This difference was particularly pronounced in the HSPC dataset, highlighting the importance of the presence barcode for reliable enhancer quantification in primary cells, where detection rates for low-activity enhancers may otherwise be limited.

Together, these results demonstrate that sc-lentiMPRA yields robust and reproducible enhancer activity measurements at single-cell resolution in both established cell lines and primary cell cultures. By combining lentiviral delivery with direct guide capture and targeted transcriptome amplification on the 10x platform, our approach provides a scalable framework for assaying enhancer activity at the single-cell level that is broadly applicable to cell lines and primary cells.

### Power analysis of sc-lentiMPRA

To guide the design of larger-scale screens, we quantified how cell numbers affect the statistical power of sc-lentiMPRA to detect activating and repressive CREs. For this, we first calculated the average activity levels of two control groups in K562 cells: 11 random DNA sequences, 11 highly active synthetic promoter sequences (JeT promoter [34]) and 10 sequences containing 4-6 repeats of Runx1 motifs, that we previously identified as strong repressors in K562 cells [16] (Additional File 1: Fig. S8a). Notably, the latter repress below the baseline activity of the random DNA sequences. To assess detection power as a function of cell number (defined as presence barcode positive cells), we downsampled the number of observations for all sequences and performed 100 pairwise Wilcoxon rank-sum tests between activator, random, and repressor sequence groups. Our results indicate that as few as 10–20 single-cell measurements are sufficient to detect significantly increased activity of the strong activator relative to random DNA controls (Fig. S8b). In contrast, we required at least 40 single-cell measurements to detect repression relative to random DNA controls (Additional File 1: Fig. S8b). Repeating this analysis in HSPCs yielded similar results for activators, with significant activity differences detectable using 10-20 single-cell observations (Fig. S8c,d). By contrast, repressive sequences in HSPCs were not robustly identified at the cell numbers tested. In summary, our sc-lentiMPRA assay enables sensitive detection of highly active enhancers with as few as 10–20 single-cell measurements per cell state and CRE.

### sc-lentiMPRA characterizes cell type specificity of synthetic enhancers at high resolution

We next applied sc-lentiMPRA to characterize synthetic enhancers across a pan-myeloid differentiation landscape of mouse HSPCs. To this end, we designed library Alpha, containing 84 synthetic sequences to explore cell type specificity of minimalist enhancer architectures containing 2-6 motifs of seven hematopoietic TFs (Cebpa, Gata1, Gata2, Spi1, Fli1, Gfi1b, and Trp53) and three TF motif pairs (Cebpa-Gata2, Fli1-Spi1, Gata1-Gfi1b) at single-cell resolution (Fig. 2a). These TFs and TF combinations were chosen based on bulk MPRA data of synthetic enhancers [16]: Enhancers containing the selected TF motifs or TF motif combinations displayed a large dynamic range (including repressors, weak and strong activators). Synthetic enhancers were designed by placing varying numbers of TF motifs into random DNA sequences, as described (ref. [16] and methods). The background DNA sequences were filtered to remove strong instances of TF motifs, without aspiring to create a “biochemically inert” background. In our previous bulk MPRA study, we found that random background DNA accounted for approximately 42–56% of the variance in enhancer activity [16]. In total, library Alpha measured 45 single-motif, 24 motif-pair and 15 random background sequences in a total of 66,582 cells. We also performed a second sc-lentiMPRA screen of 72 enhancers with Cebpa binding sites, termed library Beta and described below.

**Fig. 2:**
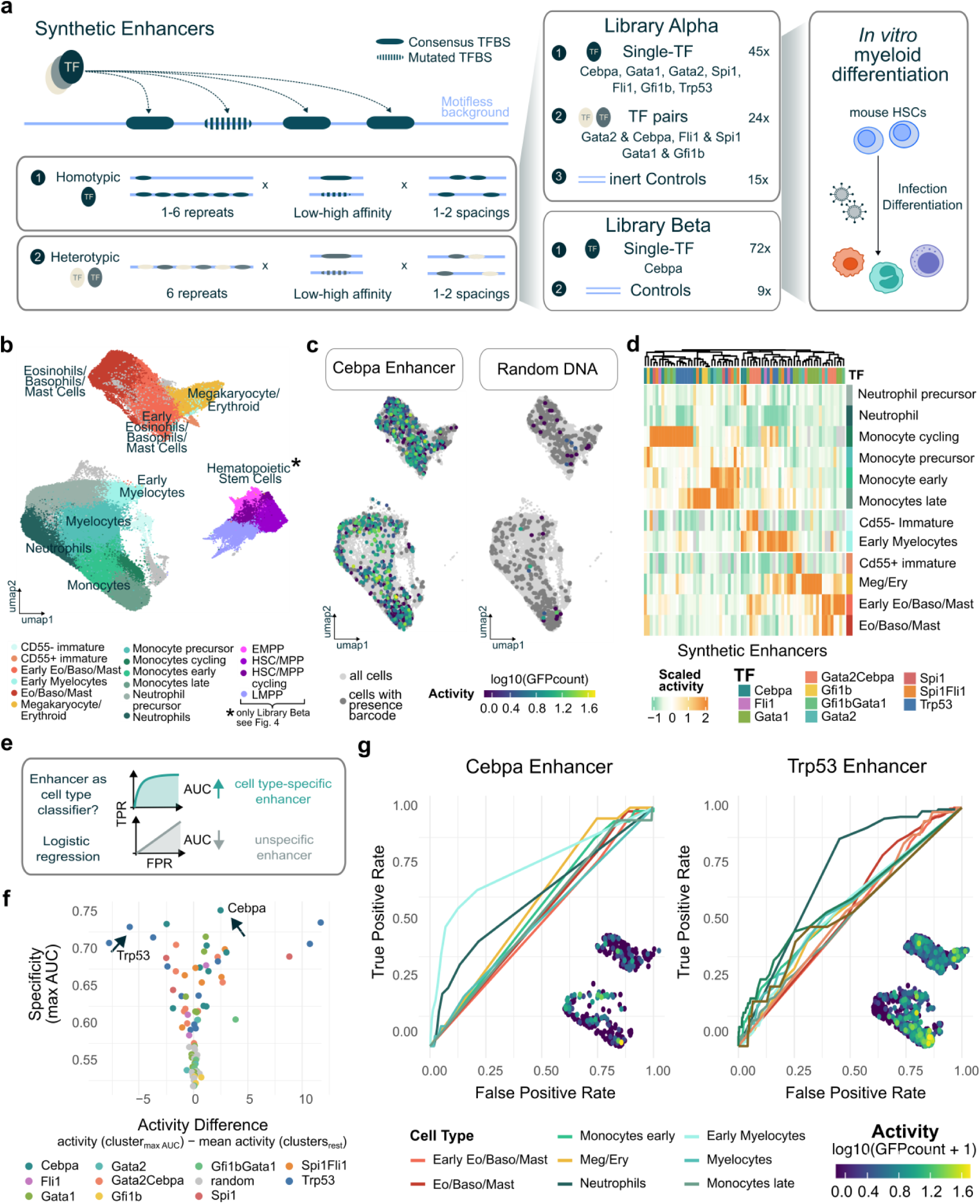
Synthetic enhancer library shows cell type-specific regulatory activity during HSPC differentiation. **a** Synthetic enhancers were generated by embedding consensus and progressively mutated motifs in a motifless random background sequence. Library Alpha comprised single-TF enhancers for seven hematopoietic TFs (Cebpa, Gata1, Gata2, Spi1, Fli1, Gfi1b, Trp53), TF pairs enhancers combining selected TF pairs (Cebpa-Gata2, Fli1-Spi1, Gata1-Gfi1b), and random DNA control sequences. This library contained 84 candidate enhancers and was assayed during *ex vivo* differentiation of mouse HSCs into myeloid progenitors. Library Beta included 72 synthetic enhancer sequences and 9 random DNA control sequences. The synthetic enhancers contained Cebpa motifs of 1-6 instances and 5 different affinities. In contrast to library Alpha, this library was additionally assayed in hematopoietic stem cells maintained in stem cell expansion cultures [23]. **b** Integrated UMAP projection of single-cell transcriptomes from the *ex vivo* HSPC differentiation culture of Library Alpha and Library Beta screen, profiled using a custom 287-gene targeted panel (Alpha: n = 66,582 cells; Beta: n = 129,787 cells). **c** Enhancer activity overlaid on the UMAP, for a synthetic Cebpa enhancer in comparison to a random DNA control sequence. **d** Clustered heatmap of single-cell enhancer activity for library Alpha, with activity values scaled per enhancer across cell states to highlight relative cell type-specificity patterns. **e** Cell type specificity of each enhancer was quantified using a logistic regression classifier trained on that enhancer’s activity, with performance summarize by ROC AUC; higher AUC indicates greater specificity for the target cell type. **f** Summary of enhancer specificity and activity difference across all 84 synthetic enhancers. Specificity is defined as the maximum AUC achieved by each enhancer across cell types, while activity difference indicates whether the enhancer is active or inactive in the corresponding cell type compared to the rest. Arrows highlight the enhancers shown in g. **g** Representative Cebpa-, and Trp53-containing enhancers illustrating cell type-specific enhancers.

The CRE library Alpha was lentivirally delivered to *ex vivo* HSC expansion cultures [23] and subsequently cultivated for six days under conditions promoting pan-myeloid differentiation [16,35]. We thereby obtained early, late and cycling Monocytes, Myelocytes, Early Myelocytes, Neutrophils and Neutrophil precursors, early Eosinophils/Basophils/Mast cells, Eosinophils/Basophils/Mast cells and a population of Megakaryocyte/Erythroid cells, which we annotated based on marker gene expression (Figure 2b, Additional File 1: Fig. S9). In the Library Beta experiment, we additionally assayed HSCs in stem cell expansion cultures [23]. These cultures contained Erythroid Multipotent Progenitors (EMPPs), cycling and non-cycling HSC and Multipotent Progenitors (MPPs), and Lympho-myeloid multipotent progenitors (LMPPs).

Having defined these cellular states, we next visualized enhancer activity across the pan-myeloid differentiation landscape. Representative UMAP projections illustrate a Cebpa enhancer with broad activity in myeloid cells, and no activity in Megakaryocyte/Erythroid cells, as well as a random DNA control in which only presence barcodes are detected in the majority of cells (Fig. 2c). These observations illustrate that sc-lentiMPRA can resolve differential activity in a complex primary system. Construct detection was broad and consistent across diverse hematopoietic states and not restricted to specific lineages (Additional File 1: Fig. S10). Each candidate CRE was detected in a median/mean of 34/56 cells per cell state (Alpha: 25/35, Beta: 35/59, Additional File 1: Fig. S10c) corresponding to a median coverage of 485 and 761 cells, respectively, per enhancer across all cell states.

We then averaged activity values per sequence across all cells of each cell state and visualized scaled activity values per enhancer in a clustered heatmap (Fig. 2d). The clustering of cell types based on enhancer activities largely recapitulated transcriptomes and known developmental relationships between cell types (Additional File 1: Fig. S11). For example, Cebpa constructs showed decreased activity in erythroid precursors and activity in all, or subsets of, myeloid cells, consistent with Cebpa’s established role in myeloid differentiation. In contrast, the Gata2-Cebpa heterotypic combinations exhibited strongest activation in early progenitor states, illustrating how TF combinations and non-linear interactions between TFs can shift enhancer specificity [16].

Next, we sought a systematic and quantitative metric for cell type specificity of enhancer activity that is robust to differing cell numbers across lineages. We therefore used logistic regression to test, for each enhancer and cell type, how well enhancer activity predicted cell type identity. For each enhancer–cell type pair, we summarized classification performance by the area under the receiver operating characteristic curve (AUC), which measures how well enhancer activity separates cells of one lineage from all others (Fig. 2e). An AUC of 0.5 indicates no separability, whereas values approaching 1.0 reflect strong and consistent enrichment of enhancer activity in a given cell type. In addition to the specificity, we additionally characterized activity difference, thereby distinguishing enhancers that are selectively active in one lineage from those that are broadly active but selectively inactive in a particular lineage (Fig. 2f).

This analysis revealed that several synthetic enhancers, including enhancers containing Trp53 and Cebpa motifs (Fig. 2f), exhibited larger cell type specificity, whereas random DNA controls displayed largely uniform activity across cell types (Additional File 1: Fig. S12). For example, an enhancer containing Cebpa binding sites showed highly specific activity to early myelocytes (Fig. 2g). Interestingly, a Trp53-driven enhancer was classified as specific to neutrophils, not due to high activity, but due to selective inactivity in this lineage compared to others, highlighting the utility of this approach in capturing both positive and negative specificity.

Together, this analysis demonstrates the capacity of sc-lentiMPRA to quantify cell state specificity of candidate enhancers. Notably, such quantitative and comparative analysis across cells is difficult to achieve in bulk MPRA, where each population is processed separately and baseline levels are difficult to assess and compare. In contrast, our single-cell approach enables direct quantification of enhancer activity and specificity in biologically complex systems within a single experiment.

### Low, but not high, affinity Trp53 motifs create enhancers sensitive to Trp53 levels

The combination of sc-lentiMPRA and synthetic enhancer libraries uniquely enables control over enhancer design alongside quantification of activity across a continuum of diverse cell states with varying TF expression. It therefore can provide data to systematically dissect the relationship between enhancer sequence, TF expression, and resulting enhancer activity. To illustrate this concept, we first focused on 12 enhancers within Library Alpha with binding sites for tetrameric Trp53 placed in random DNA. These included enhancers with 2, 4 and 6 Trp53 motifs of very high, high, medium or low affinity (Fig. 3a). Here, very high affinity sites are defined as the most likely sequence under Trp53’s PWM, whereas low affinity sites are randomly sampled from the PWM constrained to lower likelihoods [16] (see Methods). All instances of low affinity binding sites therefore were different from each other.

**Fig. 3.**
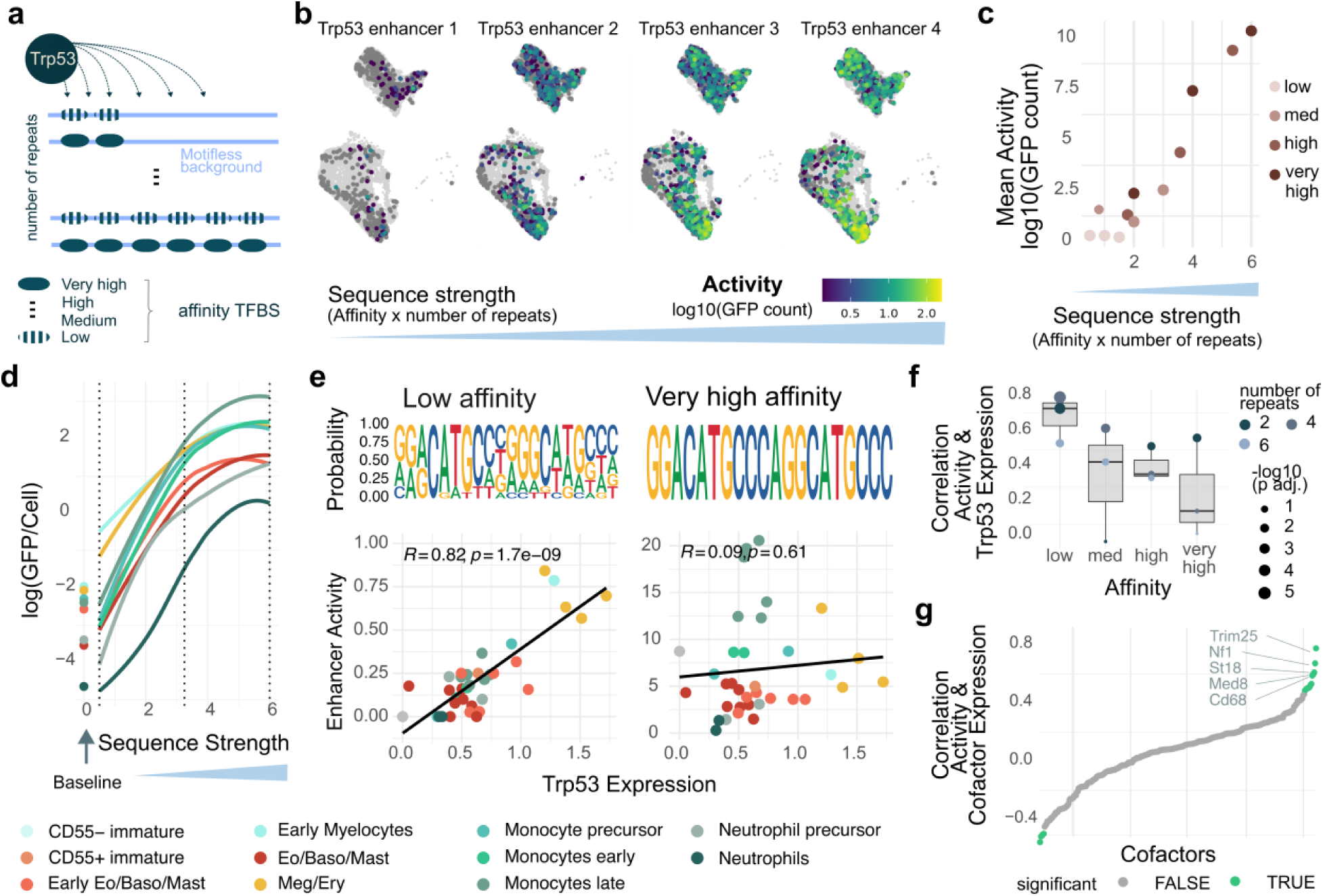
Determinants of Tpr53-containing enhancer activity across hematopoietic cell states. **a** Sublibrary of 12 synthetic enhancers comprising 2, 4, or 6 Trp53 binding sites embedded in random, motifless DNA. For each motif number, binding sites spanning very high, high, medium, and low affinity were sampled according to Trp53 PWM likelihood. **b** Single-cell activities of four example Trp53 enhancer sequences displayed on the UMAP from figure 2b. **c** Relationship between sequence strength, defined as the product of affinity and repeat number, and mean enhancer activity aggregated across all cell states. Each point represents one enhancer design. Activity increases approximately linearly with sequence strength. **d** Enhancer activity stratified by cell state. Points indicate baseline activity of background random DNA control enhancers. **e** Correlation analysis across cell states reveals strong association between Trp53 enhancer activity and Trp53 expression for low-affinity enhancers, but no association for highest-affinity enhancers. Sequence logos indicate distribution of bases in the motifs that were actually used in the synthetic enhancers with low and high affinity motifs, respectively. **f** Summary of activity-expression correlations across enhancer affinities, with progressively decreasing correlation with decreasing motif affinities. **g** Correlation of Trp53 enhancer activity with expression of transcriptional coregulators. Multiple factors remain significant after multiple-testing correction and show correlations exceeding that of Trp53 expression itself (False Discovery rate < 0.05).

Across cell states, we observed a clear positive relationship between enhancer activity and the number and affinity of Trp53 binding sites (Fig. 3b,c). Stratification by cell state revealed cell state-specific saturation behavior, with plateaus at different activity levels across states (Fig. 3d). Notably, cell types demonstrated distinct dynamic ranges of activity across different enhancer strengths: For example, in Meg/Ery progenitors enhancers with low-affinity binding sites displayed relatively high activity, whereas activity of enhancers with many high-affinity sites were highest in late monocytes.

To examine the relationship between Trp53 enhancer activity and Trp53 expression more systematically, we clustered cells at higher resolution such that most enhancer sequences were represented by a median of at least 10 cells per cluster, in line with results from the previous power analysis (Additional File 1: Fig. S13a,b). Interestingly, enhancer activity correlated strongly with Trp53 expression at low motif affinities (R = 0.82), with progressively weaker correlations at intermediate affinities, and no significant correlation for the highest-affinity enhancers (Fig. 3e,f). These effects go beyond spurious correlations between baseline activities of the random DNA controls and Trp53 enhancer activity (F-test: p = 1.2 × 10^−6^, Additional File 1: Fig. S13c,d), and indicate a strong association between Trp53 expression and enhancer activity specifically at low motif affinities. Together, these findings are consistent with a model in which Trp53 binding becomes saturated at high-affinity binding sites, such that further increases in Trp53 abundance do not result in proportional increases in enhancer activity. Under these conditions, enhancer output may instead be constrained by additional regulatory factors, including cofactor availability.

To explore whether cofactor availability may contribute to the observed cell type-specific enhancer activity levels, we examined the correlation between Trp53 enhancer activity and the expression of a curated set of transcriptional coregulators selected from literature [36,37]. To estimate expression of these factors, sc-lentiMPRA cells were projected onto an independent 5′ 10x whole transcriptome dataset generated from the same *in vitro* culture system (see Methods). Several candidate cofactors exhibited correlations with enhancer activity that exceeded those observed for Trp53 itself and remained significant after correction for multiple testing (Fig. 3g). Notably, these candidates tended to have a higher mean interaction score with Trp53 in the STRING database compared to non-significant or negatively correlated factors (Additional file 1: Fig. S13e). Among these candidates, Trim25 exhibited the strongest and most significant correlation with Trp53 enhancer activity (R = 0.78, adjusted P = 6 × 10^−7^, Additional File 1: Fig. S13f), and has previously been implicated in the regulation of Trp53 stability, consistent with a potential functional link to Trp53-dependent enhancer regulation [38,39].

In summary, sc-lentiMPRA of synthetic enhancers revealed that Trp53 enhancers containing low, but not high affinity binding sites, show activity levels that scale with Trp53 expression across cell states, and that co-factor abundance may impact cell type-specific enhancer activity. Such differences in regulatory control may enable distinct transcriptional responses to Trp53 activation across hematopoietic lineages.

### A complex relationship between Cebpa expression, enhancer composition, and activity

Finally, we performed a second sc-lentiMPRA screen comprising 72 enhancers containing varying numbers of Cebpa binding sites with different affinities, together with 9 control sequences (5 random DNA controls, 4 strong JeT promoter sequences [34]), assayed across 129,787 cells after quality control filtering (Fig. 4a). As before, motifs of different affinities were obtained by sampling according to their probability under a Cebpa position weight matrix derived from the CIS-BP database (M00994_2.00-Cebpa). Due to the low complexity of the Cebpa PWM, this resulted in a single DNA sequence for most target affinities (Fig. 4b). In contrast to the Library Alpha screen, we extended the experimental HSPC culture protocol to additionally sample cells cultivated in stem cell expansion medium ([23] and see Methods), before the initiation of differentiation. This enabled us to additionally investigate transcriptional activity in HSCs, EMPPs and LMPPs (see also Fig. 2b).

**Fig. 4.**
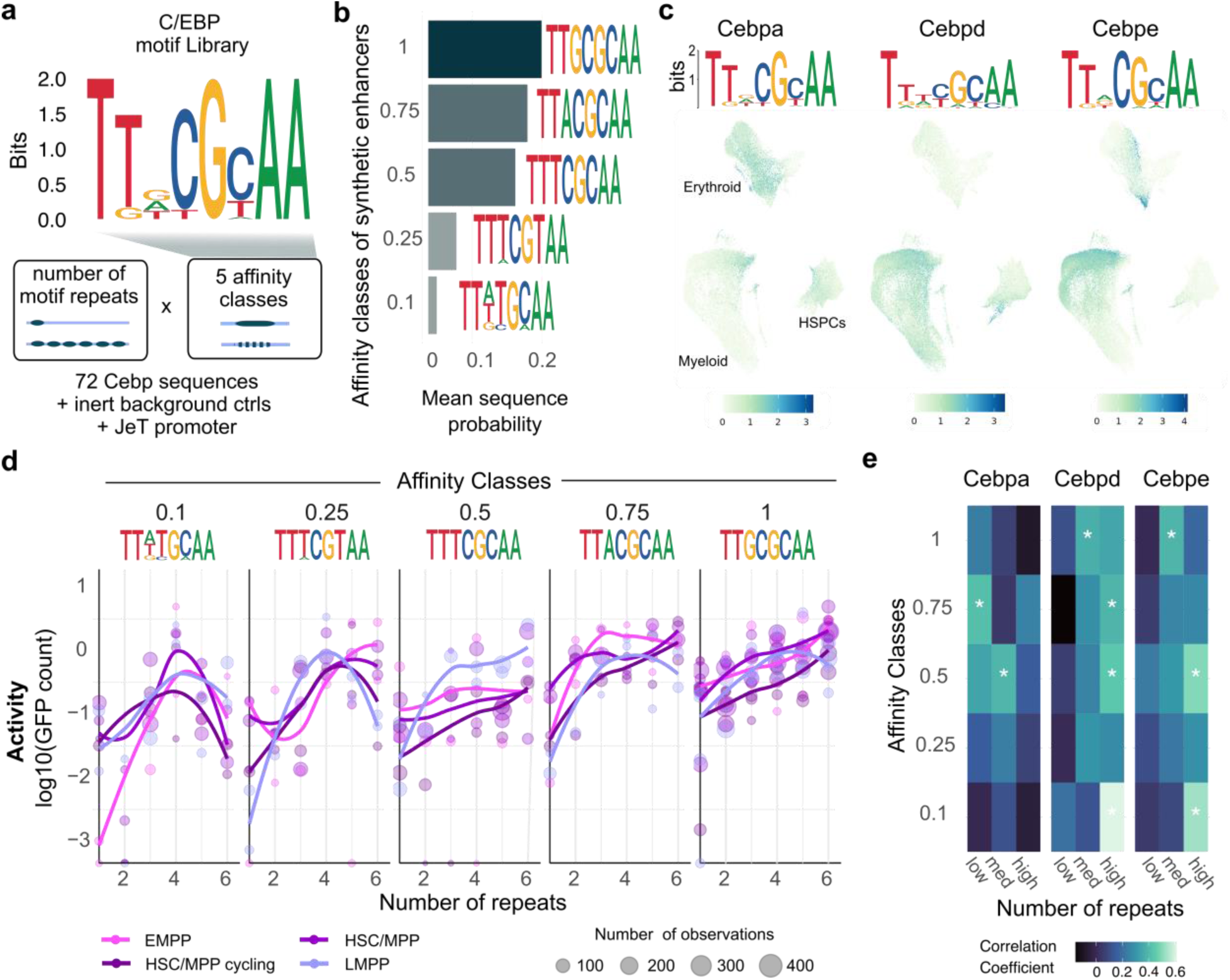
Synthetic C/EBP motif library Beta reveals non-linear enhancer regulation during HSPC differentiation. **a** Library Beta comprises 72 sequences containing 1-6 C/EBP motif repeats spanning five target affinities, together with five random DNA control sequences and 4 strong promoter sequences (JeT promoter). **b** Motif instances were sampled from the Cebpa PWM derived from CIS-BP (M00994_2.00-Cebpa) to achieve defined affinities (Methods); here, 5 different target affinities were used. Sequence logos depict sequences obtained for each target affinity. Bar chart on the right depicts the mean probability of the generated motifs, given the PWM, stratified by affinity class (0.1 −1). **c** UMAP projection of hematopoietic differentiation (see Fig. 2b annotation), with cells colored by expression levels of Cebpa, Cebpd and Cebpe. PWMs for the three TFs from CIS-BP are also shown. **d** Enhancer activity as a function of Cebpa motif number, stratified by activity classes and cell state (HSCs, EMPPs and LMPPs). **e** Correlation coefficients of Cebpa enhancer activity and expression of Cebpa, Cebpd and Cebpe, shown for 9the 5 different affinity classes and for low (1-2), medium (3-4) and high (5-6) motif repeat numbers (p-value < 0.05).

When relating motif number to enhancer activity within single cell states, we observed non-linear behavior, in contrast to the linear relationship between sequence strength and activity observed for Trp53. Enhancers containing high-affinity motif variants showed increases in activity upon the addition of further motif copies, whereas low-affinity motif variants exhibited pronounced non-linear behavior, with maximal activation at intermediate motif numbers. This behavior was most clear in HSCs and early progenitors (Fig. 4d), but also apparent in other cell states (Additional File 1: Fig. S14), and is in line with data from a bulk MPRA screen, where the relationship between motif number and enhancer activity was equally found to be non-monotonous across 270 Cebpa enhancers [16]. This observation raises the question if non-linear relationships also govern the relationship between TF expression and enhancer activity.

Of note, Cebpa, Cebpd and Cebpe are all expressed in our biological system and share highly similar binding sites (Figure 4c), suggesting that multiple CCAAT/enhancer binding protein (C/EBP) family members may contribute to the observed enhancer activity. We therefore correlated enhancer activity with Cebpa, Cebpd, and Cebpe expression separately for sets of enhancers defined by motif affinity and repeat number, across cell states (n = 45 subclusters) (Additional File 1: Fig. S15a,b). We found that some enhancer designs were correlated with Cebpa, others with Cebpd and Cebpe, and others with none of the three factors (Figure 4e). This finding illustrates that even minor changes in motif sequence and number alter preference for different members of the same TF subfamily, in a manner that is not readily apparent from the TF’s PWM (see also Figure 4c). To investigate possible non-linear relationships, we compared models describing enhancer activity as a linear or non-linear function of Cebpa, Cebpd and Cebpe expression (Methods). We found that enhancers with medium-affinity motifs displayed evidence for a non-linear relationship between TF expression and activity (Additional File1: Fig. S15c,d).

Together, these results indicate that the activity of Cebp motif-containing enhancers is shaped by complex non-linear relationships and patterns consistent with competition between related TF family members, as well as additional regulatory complexities. They further illustrate how subtle changes in TF motif sequence and number result in different responses to several similar TFs.

## Discussion

Cell state-specific activity of enhancers is of large interest in contexts such as the systematic dissection of regulatory logic,the interpretation of non-coding genetic variants,or biotechnological applications, including cell fate engineering and gene therapy. These efforts require methods to characterize natural and synthetic enhancers in relevant primary cell models, ranging from organoids to hematopoietic stem cell cultures. To date, however, most work has relied on bulk MPRA measurements, which average across heterogeneous populations or continuous differentiation trajectories, and therefore obscure cell state-specific CRE activity. This limitation has motivated the integration of MPRAs with single-cell readouts.

Existing single-cell MPRA delivery strategies are not compatible with most primary cell systems: On one side, we showed that methods without an independently expressed detection barcode may suffer from compromised activity quantification. On the other, one system, scQers, represents an important advance by incorporating such a detection barcode into a circular RNA, offering high recovery rates [17]. However, scQers relies on piggyBac integration and lipofection, limiting its compatibility with most primary cell systems. An adaptation of scQers to lentiviral delivery is difficulted by the long distance between barcode and CRE in that system, which would increase lentiviral recombination rates and consequently reduce statistical power [33,40]. In contrast, our sc-lentiMPRA is optimized for lentiviral delivery, maintains low recombination and barcode-swapping rates due to the close proximity of barcodes and CREs, and achieves robust detection barcode recovery. This design overcomes key technical constraints of primary cell assays and provides a practical solution for reliable detection of barcode–based CRE activity quantification at single-cell resolution. By using targeted sequencing, sc-lentiMPRA achieves high on-target rates and robust detection power while substantially reducing sequencing costs.

Unlike previous work in the scMPRA context, we here focus specifically on the characterization of fully synthetic, minimalist enhancers at single-cell resolution. Minimalist enhancer constructs, composed of random DNA and TF binding sites, have emerged as a powerful tool to dissect principles of gene regulation [16,21,41–45]. Here, we demonstrate that a single-cell characterization of such enhancers can systematically dissect the relationship between enhancer sequence, TF expression levels, and enhancer activity by correlating TF expression and enhancer activity. This approach is complementary to perturbation based approaches [36,46], which may suffer from indirect effects of perturbation on cell state that are not explicitly measured. Most notably, our results suggest that motif affinity can critically shape how enhancer activity responds to TF levels.

In the example of Trp53, we propose a model in which low-affinity motifs function as dosage-sensitive enhancer elements, whose activity scales with TF concentration, whereas high-affinity motifs may be bound by Trp53 to saturation and thus respond less to changes in TF levels. Similar observations have been made for the TF TWIST1, using tight dosage control with a dTAG system [47]. Our data further suggests that activity of enhancers with high-affinity motifs may instead be governed by additional constraints such as cofactor availability or chromatin context. Consistent with this model, low-affinity binding sites have emerged as key determinants of regulatory precision, enabling context-dependent and spatially restricted enhancer activity rather than maximal transcriptional output [48–50].

The Cebpa examples highlight additional layers of complexity. On one side, it demonstrates that enhancers with very similar motif sequences or numbers can differ in correlation with the expression of related TFs. On the other side, the same low affinity Cebpa motifs were most active at intermediate repeat number. A potential explanation for this behavior is “duality”, a phenomenon whereby a TF can act as an activator at some motif number and as a repressor under others [16,51]. Like many TFs, Cebpa contains repressive and activating domains [52–54], that can incoherently affect polymerase activity [51]. Additionally, the single-cell nature of our data allows us to demonstrate that for some motif variants, changes in Cebpa expression levels are associated with non-linear changes in enhancer activity. Such effects may underlie strongly dosage-dependent, non-linear responses to TF dose observed in various cellular systems [55–57]. Together, these observations underscore that enhancer function emerges from the combined effects of motif affinity, TF dosage, and combinatorial TF interactions.

In summary, sc-lentiMPRA is a powerful lentivirus-based method for investigating enhancer activity and regulatory dynamics at the level of individual cells and is applicable to various biological systems including primary cells. We demonstrate that the single-cell profiling of minimal, designed enhancers enables detailed studies on the relationship between enhancer sequence, TF expression and enhancer activity during cellular differentiation.

### Limitations

A limitation of our system is that efficient barcode capture requires stable Cas9 or dCas9 expression in the cells of interest to promote gRNA-barcode stability. Successful CRISPR and Perturb-seq screens illustrate that this requirement can be achieved also in human primary cells [58,59]. A second limitation is that the current implementation of sc-lentiMPRA relies on a specific commercial platform (10x Genomics).

## Data availability

Seurat objects containing raw and normalized gene expression, presence barcode and quantification barcode calls are available at https://doi.org/10.6084/m9.figshare.31420724

## Acknowledgements

We thank Rosa Martínez Corral for discussion throughout the project. We thank the CRG facilities for Genomics and Flow Cytometry, as well as the PRBB animal house, for experimental support. This study was financed by grants from the Spanish Ministry of Science, Innovation and Universities (MICIU/AEI /10.13039/501100011033) with cofunding from the European Union NextGenerationEU/PRTR/FSE+ (grant PID2019-108082GA-I00 to L.V. and pre-doctoral fellowship PRE2021-097675 to J.R.) and the European Union’s Horizon Europe under the grant agreement No 101041399 (ERC-StG AI4SYN to L.V.) The authors acknowledge support of the Spanish Ministry of Science and Innovation to the EMBL partnership, the Centro de Excelencia Severo Ochoa and the CERCA Programme / Generalitat de Catalunya.

## Author contributions

JR, RF and LV conceived of the study. JR developed sc-lentiMPRA and performed experiments, with support by ABM, RF, and CST. JR analyzed the data and wrote the manuscript. LV supervised work. RF and JB provided guidance. All authors commented on the manuscript.

## Methods

### Construction of the sc-lentiMPRA vector

To generate the scMPRA backbone vector, we used the lentiviral reporter plasmid pLS-SceI (Addgene #137725), which contains multiple restriction sites upstream of an EGFP reporter gene [24]. The plasmid was digested with SbfI, and the hu6 promoter (amplified from CROPseq-PURO-Gfp F+E [22]) was inserted by Gibson assembly to create the scMPRA backbone. The sc-lentiMPRA vector is available upon request.

### Design of Synthetic Enhancer Libraries PoC, Alpha, Beta

Our library design approach and relevant scripts are described in [16]. In brief, each synthetic enhancer sequence was created from three components: TFBSs, spacers between the TFBSs, and filler sequences to standardize length. To avoid systematic biases, we used random DNA as spacer and filler sequences, rather than a fixed background sequence. Thereby we avoided that cryptic motifs in a single background sequence might introduce systematic biases. Background DNA was filtered with FIMO [60] to avoid motifs of 38 hematopoietic TFs. This does not avoid the occasional appearance of information-poor potential TF sites, such as GATA.

PWMs were obtained from HOCOMOCO v11 [61], JASPAR 2020 [62], and CIS-BP v2.0 [63], motifs were clustered, and representative centroid motifs were selected, as described [16]. To obtain mpotif instances of different affinity, 100,000 motif instances were drawn from the corresponding PWM and ranked by their likelihood given the PWM. Affinities were then defined as percentiles on the resulting rank-ordered motif list, where the 100^th^ percentile represents the most likely motif given the PWM. To select motifs of a desired strength, motifs were then sampled within a ±2.5% range around the target affinity percentile. This approach generated multiple motif instances of similar affinity. The target affinities used for the different libraries are specified below. Finally, each sequence was flanked on both sides by 15 bp lentiMPRA adaptor sequences for cloning [24].

*Library PoC* contained a total of 432 elements previously tested in bulk MPRA experiments [16], including 348 single-TF sequences, 62 endogenous enhancer sequences, and 22 control sequences. Control sequences consisted of 11 random background sequences filtered for TFBS as described above, and 11 highly active sequences of the JeT promoter [34].

*Library Alpha* contained 84 synthetic sequences containing motifs for seven hematopoietic TFs (Cebpa, Gata1, Gata2, Spi1, Fli1, Gfi1b, and Trp53) and three motif pairs (Cebpa-Gata2, Fli1-Spi1, Gata1-Gfi1b). These motifs and motif pairs were chosen because they displayed strong repressive or activating activity in bulk MPRA screens, and specificity to flow-sorted populations. For Fli1, Gata2, Gfi1b and Spi1, we used enhancers with 6 repeats of high affinity motifs (89^th^ or 100^th^ percentile of likelihood-ranked PWM samples), and created three enhancers differing in background sequence, for a total of 3 enhancers per TF. For Cebpa, we used enhancers with 6 repeats of low, mid and high affinity motifs (25^th^, 50^th^ and 89^th^ percentile of likelihood-ranked PWM samples), and created three enhancers differing in background sequence for each affinity, for a total of 9 enhancers. For Gata1, we used 1, 3, 4 or 6 repeats of mid and high affinity motifs (50^th^, 75^th^, 89^th^ percentile), for a total of 12 enhancers. For Trp53, we used 2, 4 or 6 repeats of low, mid and high affinity motifs (25^th^, 50^th^, 89^th^ and 100^th^ percentile), for a total of 12 enhancers. For the TF combinations, we used 3 repeats of high affinity motif pairs (90^th^ percentile), in alternating order, and explored 2-3 spacing options per pair, for a total of 24 enhancers. Spacing and orientation of motifs were chosen based on bulk MPRA results [16]; of note, spacing and orientation did not have a substantial impact on TF activity for any of the single TFs selected, but spacing played a role for the TF pairs. Finally, we included 15 random background sequences.

*Library Beta.* Library Beta included enhancers containing 1-6 Cebpa motifs of different affinities (10^th^, 25^th^, 50^th^, 75^th^, 89^th^ and 100^th^ percentile of likelihood-ranked PWM samples), for a total of 36 site number/motif affinity combinations. Spacing between Cebpa sites was fixed at 10 bases and motifs were placed in forward orientiation, since our previous bulk screen of 270 Cebpa-motif containing sequences had identified no substantial effect of motif spacing/orientation on activity [16]. We generated two different sequences for each combination of site number and motif affinity, differing in background. Additionally, we included 9 controls (5 random DNA controls, 4 strong JeT promoter sequences [34]).

All CRE sequences are included in Supplementary table 3, and a more detailed annotation alongside activity measurements is available on figshare (see data availability statement).

### Cloning of enhancer libraries

For a visual overview of enhancer library cloning, see figure S2a. DNA libraries were ordered from Twist Bioscience. In two consecutive PCRs, adapted from lentiMPRA, we appended necessary sequence elements: in the first PCR, a hu6 promoter and tracrRNA were added on one side of the CRE, and part of a minimal promoter on the 3’side. In the second PCR the gRNA barcode was added on the 5’ side of the insert and the GFP barcode, and an EGFP homology overhang were appended on the other side. The first PCR was performed with primers #37 and #212, and the second with primers #242 and #243.

To optimize amplification, an initial qPCR with primers #37 and #212 was used to determine the minimal cycle number for the first-round PCR. To prevent PCR jackpotting, multiple parallel reactions were performed, yielding at least 100 ng of first-round product (9–14 cycles; 14 for PoC, 11 for Alpha and 9 for Beta). Second-round PCRs were run for 4 cycles set up in multiple PCR reactions as for the first PCR. The second PCR step introduced both the guide and GFP barcodes as well as the Gibson overhangs.

The scMPRA backbone vector was linearized with AgeI-HF and Esp3I, and purified insert DNA from the second-round PCR was assembled into the backbone using Gibson assembly (NEBuilder HiFi DNA Assembly). To eliminate residual undigested backbone, an I-SceI digest was performed after assembly. The resulting recombinant plasmids were purified, quantified, and subsequently used for bacterial electroporation. NEB10B electrocompetent cells were pulsed once at a voltage of 2.0 kV, 200 ohms as resistance and 25 uF of capacitance. Bacteria were incubated in LB plates with ampicilin overnight at 37°C, colonies were collected the following day. For library Alpha we obtained 6.9*10^6^ colonies, and for library Beta we obtained 1.65*10^6^ colonies.

### sc-lentiMPRA CRE-Barcode association libraries

The fragment containing both the detection barcode (gRNA) and the quantification barcode (GFP) was PCR-amplified from the plasmid library using primers that added Illumina sequencing adapters and sample indices (see Figure S2a, primers #216_P5-pLSmP-ass-i742.FWD and #218_P7-pLSmp-ass-gfp.REV). Amplicons were gel-purified and subjected to paired-end sequencing with custom primers in the following format on an illumina iSeq sequencer: #219_Read1_Seq_CRS_ass.FWD (15 cycles), #220_Read2_Seq_CRS_ass.REV (110 cycles), #221_IndexRead1_Seq_CRS_ass.FWD (15 cycles), and #222_IndexRead2_Seq_CRS_ass.REV (10 cycles). Data was processed into tables linking detection barcode sequence, quantification barcode sequence and candidate enhancer sequence using a custom perl script.

### Lentivirus production

Lentivirus was produced in HEK239FT cells by co-transfecting the library plasmids (1.64 pM) with the packaging plasmid psPAX2 (1.3 pM) and the envelope plasmid pMD2.G (0.72 pM) (Addgene, Plasmids #12260 and #12259). Transfection was carried out for 6 h using Lipofectamine 3000 according to the manufacturer’s instructions (Invitrogen). Viral supernatant was collected 72h post-transfection, filtered using a 0.45 μm low-protein-binding filter, and lentiviral particles were concentrated by sucrose cushion ultracentrifugation (60,000 g, 1.5 h, 4 °C) [64]. For the K562 screen, the virus pellet was resuspended in RPMI medium, whereas for the HSPC infection we used StemSpan medium and stored at –70 °C. Viral titers were determined using the Lenti-X qRT-PCR Titration Kit (Takara Bio). Prior to MPRA screening, each viral preparation was validated in K562 cells and HSPCs to confirm transduction efficiency, cell viability, and GFP reporter activity.

### Mice, cell culture and lentiviral infection

To achieve gRNA stability and enable efficient gRNA capture, all screens were performed in cells expressing nuclease-dead variants of Cas9. For the K562 screens, we worked with a previously characterized clone of cells expressing dCas9-BFP-KRAB [22]. For the HSPC screens, we used primary cells derived from bone marrow of mice expressing dCas9-KRAB in the hematopoietic lineage. These mice were obtained by crossing B6J-LSL-dCas9-KRAB (obtained from the Jackson Laboratory, strain ID 033066 [65]) with Vav1-Cre mice (obtained from the lab of Thomas Graf, Jackson strain ID 035670 [66]). All procedures involving animals adhered to the pertinent regulations and guidelines of the European Union, Spain, and Catalonia. Approval and oversight for all protocols and strains of mice were granted by the Institutional Review Board and the Institutional Animal Care and Use Committee at PRBB, under protocol 11733.

HSPCs were obtained by sacrificing male and female mice of 8-20 weeks of age, and as described before [16] with minor modifications. In this specific case, we avoided the use of FBS and performed the extraction in PBS without FBS supplementation, in order to prevent premature differentiation of the cells. Briefly, lineage-negative cells were enriched with the Lineage Cell Depletion Kit (Miltenyi Biotec) and further selected for c-Kit positivity using CD117 MicroBeads (Miltenyi Biotec). The resulting Lin^−^c-Kit^+^ population was used.

For culture, freshly isolated HSPCs were plated in 24-well CellBind plates (Corning) using stem cell expansion cultures [23] consisting of F12 supplemented with HEPES (10 mM), Penicillin–Streptomycin–Glutamine (1×), ITS-X supplement (1×), and polyvinyl alcohol (1 mg/ml), further supplemented with mSCF (10 ng/ml) and TPO (100 ng/ml). Cells were maintained in a hypoxia incubator (5% O₂, 5% CO₂) at 37 °C, with medium changes every 2 days. Culture conditions were assessed on day 15 by flow cytometry to monitor the presence of Lin^−^ Sca-1^+^ c-Kit^+^ (LSK) cells. After d14, cells were grown at 37°C and 5% CO2. On day 20 of expansion culture, HSPCs were infected with lentivirus for 20 h. Specifically, we infected 24 million HSPCs for 20 hourswith a lentiviral titer between 1.6*10^10^ and 2.02*10^11^. After stopping the infection, cells recovered in Wilkinson medium for 7 days to allow stable integration and expression of gRNAs and GFP. On day 21, cells were transferred into differentiation medium, enabling differentiation across the pan-myeloid spectrum [16,35]. On day 27 post-extraction, following 6 days of differentiation, cells were collected for sc-lentiMPRA library preparation.

K562-BFP-dCas9 cells were maintained in RPMI medium supplemented with 10% FBS and 1% penicillin–streptomycin at 37 °C with 5% CO₂. Cells were infected for 20h and were allowed to recover for 7 days on the incubator prior to sc-lentiMPRA library preparation.

### Flow cytometry

We assessed differentiation progress using a previously validated flow cytometry protocol [16]. This protocol separated the differentiation cultures into seven defined cellular states: immature progenitors (CD55^−^Itga1^−^Sca1^+^), megakaryocyte-erythroid progenitors (CD55^+^c-Kit^−^CD274^−^), eosinophil (CD55^+^c-Kit^−^CD274^+^), basophil (CD55^+^c-Kit^+^), neutrophil precursors (CD55^−^Itga1^−^Sca1^−^), and early and late monocyte precursors (CD55^−^Itga1^Low^Sca1^+^ and CD55^−^Itga1^High^Sca1^+^, respectively).

HSPCs were stained with CD55-AF647, c-Kit-BV605, CD274-PE, Itga1-APC/Fire 750, and Sca-1-BV785. DAPI was used as a viability dye. The full set of markers was only used to control differentiation culture quality. To achieve a balanced representation of all cell states in the sc-lentiMPRA data, cells were then sorted into alive CD55^+^ and CD55^−^ populations on a FACSAria II cell sorter (BD Biosciences, San Jose, CA). Equal numbers of CD55^+^ and CD55^−^ cells were processed further.

### sc-lentiMPRA Library preparation

To generate gRNA, GFP, and TAP libraries, we combined and adapted previously published strategies for targeted library preparation, optimizing them for compatibility with the 10x Genomics Single Cell 5′ kit (v2) [22,30]. After FAC sorting, reverse transcription (RT) was performed using the 10x Chromium system, supplemented with guide_RT primer #46 (final concentration: 160 nM) [67] and our custom Poly-dT primer #144 (final concentration: 9.73 µM) in a 75 µl RT mix. GEM generation and RT cycling were carried out according to the manufacturer’s protocol (53 °C 45 min; 85 °C 5 min; 4 °C hold), followed by GEM–RT cleanup.

cDNA was then amplified with targeted primers to capture gRNA, the TAP genes, and GFP fragments. The reaction contained the GEM–RT cleanup product (35 µl), Partial Read1 primer #28 (10 µM, 4 µl), an HSC- or K562-specific Outer primer mix (100 µM, 2.5 µl) pooled at equimolar ratio, GFP Outer primer #63 (100 µM, 2.5 µl), Guide Add primer #206 (100 µM, 1.2 µl), and KAPA HiFi HotStart Readymix (50 µl), adjusted with nuclease-free water to 100 µl.

Following amplification, products were separated by size using SPRIselect beads, as described [67]. Larger fragments (TAP and GFP) were retained in the pellet after a 0.6× SPRIselect cleanup, while the shorter gRNA products remained in the supernatant. The pellet fraction was washed twice with 80% ethanol and eluted in 45 µl EB, providing input for TAP and GFP library preparation. The supernatant fraction was further purified using a 1.4× SPRIselect cleanup and a subsequent 2.0× SPRI cleanup, washed twice with 80% ethanol, and eluted in 45 µl H₂O. From this point, TAP/GFP and gRNA libraries were processed in parallel.

#### 1. gRNA library (from supernatant)

##### 1.1 Enrichment PCR

Enrichment PCR was performed in a 100 µL reaction containing 10 ng supernatant eluate, 2.5 µL 10 µM SI-PCR primer #14, 2.5 µL 10 µM gd_add_v4 primer #2, and KAPA HiFi HotStart Readymix (50 µL), with nuclease-free water to volume. Cycling was 95 °C for 3 min; 8 cycles of 95 °C for 20 s, 64 °C for 30 s, 72 °C for 20 s; 72 °C for 5 min; 4 °C hold. The product was purified with 1.6× SPRIselect and eluted in 20 µL (expected size ~223 bp).

##### 1.2 Index PCR (gRNAs)

Indexing was performed in 100 µL using the enrichment PCR product at 10 ng, 2.5 µL 10 µM SI-PCR primer #143 and 2.5 µL 10 µM Next_nst_guide #47, plus KAPA HiFi HotStart Readymix (50 µL) and water to volume. Cycling was 95 °C for 3 min; 10 cycles of 95 °C for 20 s, 54 °C for 30 s, 72 °C for 20 s; 72 °C for 5 min; 10 °C hold. The product was purified with 1.6× SPRIselect and eluted in 15–20 µL (expected size ~210 bp).

#### 2. TAP & GFP Libraries

##### 2.1 Inner PCR

Inner PCR was performed on a 1:1 mixture of pellet and supernatant cDNA (10 µL each). Reactions (100 µL) contained 10 ng outer product, 4 µL 10 µM Partial Read1 primer #28, 2.5 µL 100 µM Inner K562/HSC primer mix, 1.2 µL 100 µM GFP Inner primer #55, and 50 µL KAPA HiFi HotStart Readymix, with nuclease-free water to volume. Cycling was 95 °C for 3 min; 8 cycles of 98 °C for 20 s, 67 °C for 1 min, 72 °C for 1 min; 72 °C for 5 min; 4 °C hold. Products were purified with 1.5× Ampure, washed twice with 80% EtOH, and eluted in 20 µL. The purified product was used as input for GFP and TAP index PCRs.

###### 2.1.1 Index PCR (GFP)

Index PCR for GFP was performed in 100 µL using 10 ng inner product, 2.5 µL 10 µM Sergis_SI primer #267, 2.5 µL 10 µM RPI_Nxx #262, and 50 µL KAPA HiFi HotStart Readymix, with water to volume. Cycling was 95 °C for 3 min; 10 cycles of 98 °C for 20 s, 52 °C for 15 s, 72 °C for 45 s; 72 °C for 5 min; 4 °C hold. Products were purified with 1.5× Ampure and eluted in 20 µL (expected size ~260 bp).

###### 2.1.2 Index PCR (TAP)

Index PCR for TAP was performed in 100 µL using 10 ng inner product, 4 µL 10 µM 10×_SI primer #232, 2.5 µL 10 µM RPI_N7xx primer #151, and 50 µL KAPA HiFi HotStart Readymix, with water to volume. Cycling was 95 °C for 3 min; 8 cycles of 98 °C for 20 s, 67 °C for 1 min, 72 °C for 1 min; 72 °C for 5 min; 4 °C hold. Products were purified with 1.5× Ampure, washed twice with 80% EtOH, and eluted in 20 µL.

For some GFP and TAP libraries, an additional high-molecular weight peak was observed, likely corresponding to “bubble PCR” products stemming from overcycling or excess input DNA. To correct this, a reconditioning PCR was performed.

###### Reconditioning PCR (optional)

Reactions (100 µL) contained 10 ng indexed library product, 2.5 µL 10 µM P5 Illumina primer #203, 2.5 µL 10 µM P7 Illumina primer #204, and 50 µL KAPA HiFi HotStart Readymix, with water to volume. Cycling was 95 °C for 3 min; 3–5 cycles of 98 °C for 20 s, 67 °C for 1 min, 72 °C for 1 min; 72 °C for 5 min; 4 °C hold. Products were purified with 1.5× Ampure and eluted in 15-20 µL.

Libraries were quality-checked on a Bioanalyzer, pooled at the desired ratios, and sequenced using an Illumina platform. Sequencing was performed with the following cycle configuration: Index i7, 10 cycles; Index i5, 10 cycles; Read 1, 26 cycles (cell barcode and UMI); and Read 2, 50 cycles for GFP libraries or 33 cycles for guide barcode libraries.

Targeting primers for mRNA capture were designed as described [68], except that the pipeline was adapted to select primers that bind near the annotated or inferred 5′ end of transcripts. Code for primer design for 5’ assays is available at https://github.com/veltenlab/TAPseq5. Maximally informative target genes were selected from annotated transcriptome-wide single-cell data of differentiation cultures [16] as described [22]. Specifically, the selectTargetGenes() function from the TAPseq R package was used.

### Bulk MPRA screens

For the bulk MPRA screen, we sorted 12 million CD55^+^ cells and 12 million CD55^−^ cells, with two replicates each (6 million per replicate) on a FACSAria II cell sorter (BD Biosciences, San Jose, CA).MPRA library preparation was carried out following the lentiMPRA protocol [24] with minor adaptations, as described before [16].

### Lentiviral recombination quantification

Control constructs were generated by cloning the CMV promoter into the sc-lentiMPRA vector. Following bacterial transformation, eight individual colonies were picked and expanded, each carrying a unique guide-GFP barcode pair. Plasmid DNA was isolated with the QIAprep Spin Miniprep Kit (Qiagen), sequence-verified, and pooled in equal amounts to obtain a balanced mixture of all eight constructs. This pooled plasmid DNA was used for lentiviral production as described above. K562 cells were infected with the pooled virus under standard conditions, and genomic DNA was collected 72 h post infection following the bulk MPRA workflow. Briefly, cells were pelleted, lysed in RLT buffer (Qiagen) supplemented with 1% β-mercaptoethanol, and DNA was purified using the AllPrep DNA/RNA kit (Qiagen) according to the manufacturer’s instructions. Barcode pairs were recovered using a library preparation strategy analogous to the CRE-barcode association workflow for sc-lentiMPRA. The fragment spanning the gRNA and GFP barcodes was PCR-amplified from genomic DNA using primers containing Illumina adapters and sample indices (P5-pLSmP-ass-i742.FWD and P7-pLSmP-ass-gfp.REV). PCR products were gel-purified and sequenced on an Illumina iSeq platform with custom primers to capture both barcode sequences.

### Single-cell data processing and enhancer activity quantification

Raw FASTQ files were processed with the Cell Ranger count pipeline (v2.1.1, 10x Genomics), aligning the TAP library reads to either the mm10 genome (HSPCs) or GRCh38 (K562). For the gRNA and GFP libraries, we used the cellranger Feature Barcode Analysis workflow in combination with a pre-compiled list of barcodes obtained from the CRE-barcode association library for gRNA barcode and GFP barcode counting. Obtained gene expression, GFP, and gRNA count matrices were then imported into Seurat (v5.2.1) [69].

For each sample, Seurat objects were created for the gene expression, GFP, and gRNA assays. First, we performed quality control filtering based on the number of detected features and UMI counts. Normalization and variance stabilization were performed using SCTransform, with variable features restricted to a curated list of target genes. PCA was run on the top variable TAP genes, and clustering was performed using the first 12 principal components (FindNeighbors, FindClusters with resolution = 0.35). Dimensionality reduction was performed using UMAP, and clusters were annotated based on marker gene expression.

To quantify single-cell enhancer activities, we first called enhancer presence in individual cells based on gRNA UMI counts (K562: ≥3 gRNA UMIs; HSPCs: ≥2 gRNA UMIs). These thresholds were chosen as a compromise between maximizing correlation with bulk measurements, aligning the observed distributions of guide and GFP barcodes, and retaining as many single-cell measurements as possible. We then used a two-step preprocessing strategy for enhancer activity estimation: (i) call enhancer presence using the fixed gRNA UMI threshold, and (ii) use the corresponding GFP barcode count, assigned via the association library, in enhancer-positive cells as a direct measure of enhancer activity.

### Data analysis and visualization

Data was analyzed in R using default tools (e.g. logistic regression, smoothing splines) as described in the main text and figure legends. the ggplot2 [70] and pheatmap [71] were used for data visualization.

## Supplementary materials

Additional file 1: Supplementary figures and legends

Additional file 2: Primers used for scMPRA cloning, association, sequencing, and library preparation.

Additional file 3: Primer Sequences for Targeted Transcriptome Profiling

Additional file 4: Candidate enhancer sequences

