## Supplementary Figures for "Lentiviral single-cell MPRA of synthetic enhancers reveals motif affinity-based encoding of cell type specificity"

### Additional File 1

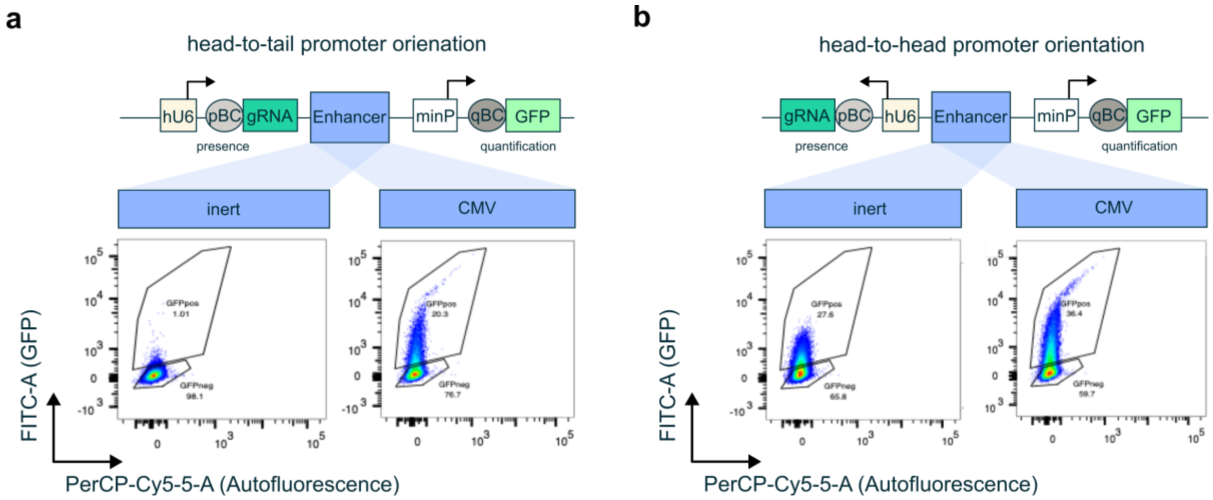


**Supplementary Fig. 1 | Vector architecture minimizes promoter cross-talk in sc-lentiMPRA.** Schematic of the sc-lentiMPRA vector design comparing head-to-tail (a) and head-to-head (b) orientations of the Pol III–driven hU6 promoter and the Pol II–driven minimal promoter. Flow cytometry analysis of GFP expression for inert and CMV enhancer constructs promoter orientations. Head-to-tail arrangement of the hU6 and minimal promoter cassettes results in reduced cross-talk and improved independence between the two transcriptional units relative to the head-to-head configuration. pBC, Presence Barcode; minP, minimal Promoter; qBC, quantification Barcode.

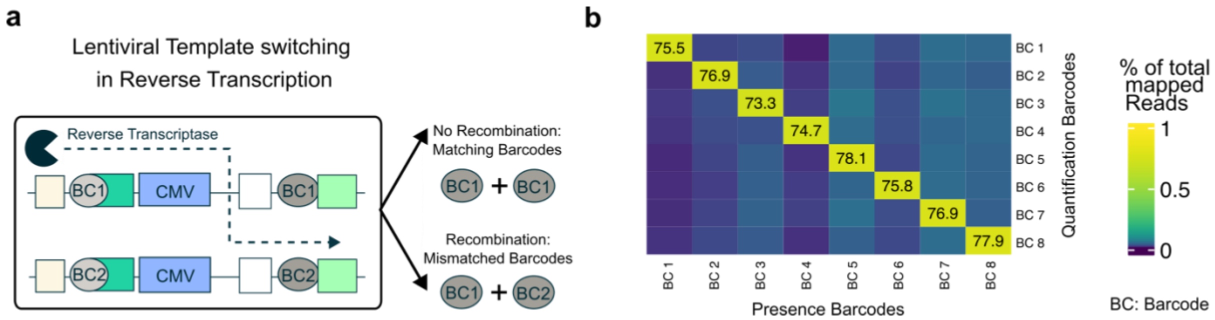


**Supplementary Fig. 2 | Vector architecture minimizes barcode swapping in sc-lentiMPRA. a** Schematic illustration of barcode swapping introduced through lentiviral template switching during lentivirus production. **b** Experimental assessment of barcode pairings to quantify lentiviral recombination in K562 cells. Eight control constructs containing only the CMV promoter with different GFp and gRNA barcodes were cloned separately, but then pooled in equal amounts for lentiviral production. The fragment spanning the gRNA and GFP barcodes was amplified from genomic DNA and sequenced. The frequency of each barcode pairing across integrated constructs for the head-to-tail configuration is visualized in the heatmap. Close physical proximity of the two barcodes within the vector backbone minimizes homologous regions and reduces lentiviral template switching. Notably, the recombination rates measured in this assay represent an overestimation of barcode swapping under library conditions. In this control setup, all constructs share an identical CMV promoter sequence, increasing the likelihood of template switching between homologous viral genomes. In contrast, in complex enhancer libraries, sequence diversity between constructs reduces the probability of productive barcode swapping events.

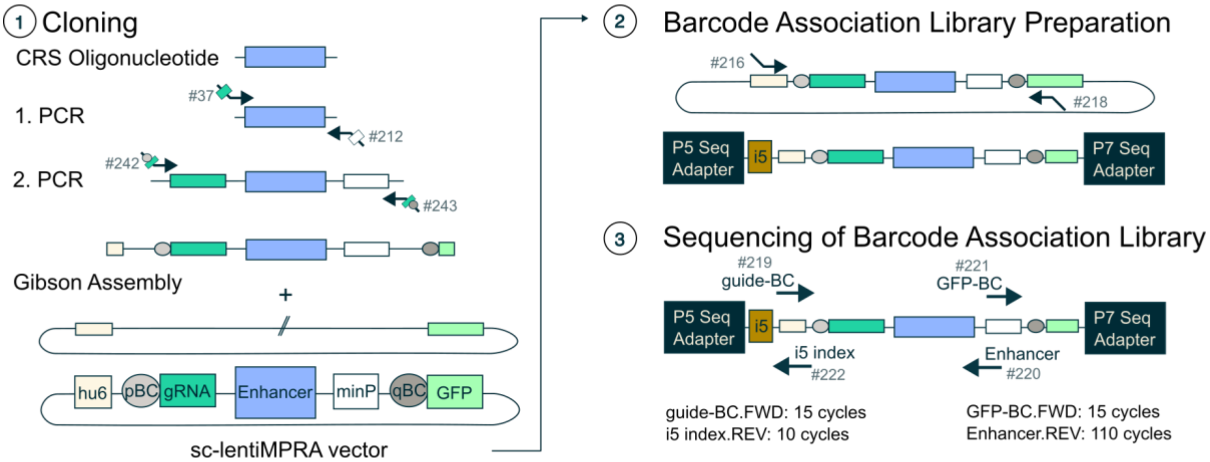


**Supplementary Fig. 3 | Cloning Procedure and Barcode Association Libraries.** The complex enhancer library in the sc-lentiMPRA vector was cloned using a two-step PCR approach that introduced the gRNA scaffold, the minimal Promoter (minP), and two barcodes: the presence barcode (pBC) and the quantification barcode (qBC) (1). To map each enhancer to its corresponding pBC-qBC combination, fragments were amplified from the plasmid library (2) and sequenced using custom primers (3).

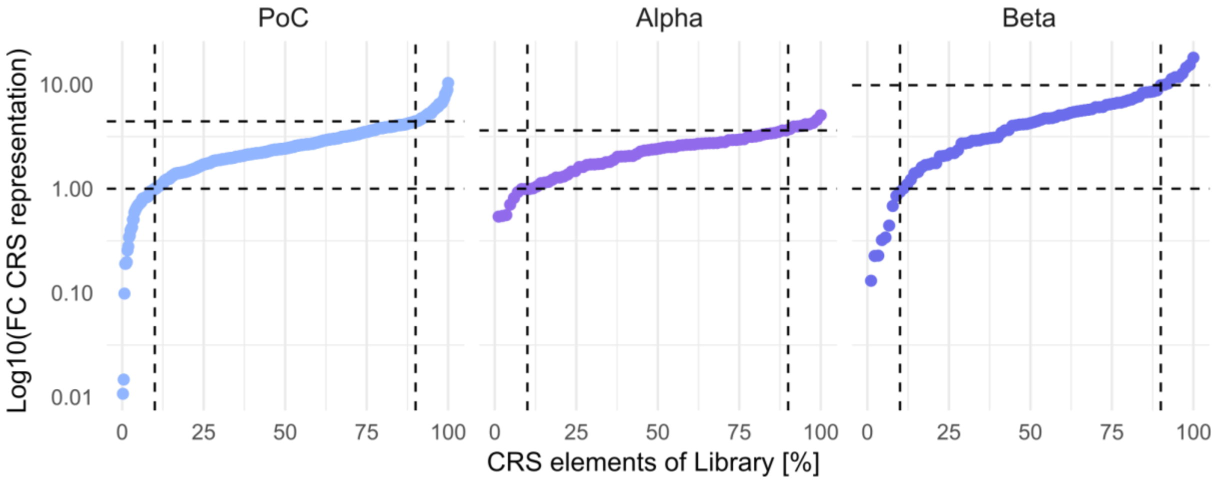


**Supplementary Fig. 4 | CRS representation in plasmid library.** Distribution of enhancer representation in plasmid library PoC, library Alpha, and library Beta after cloning. Each point indicates one element (putative enhancer) in the library and dotted lines visualize the 10$^{th}$ and 90$^{th}$ percentiles library elements.

a

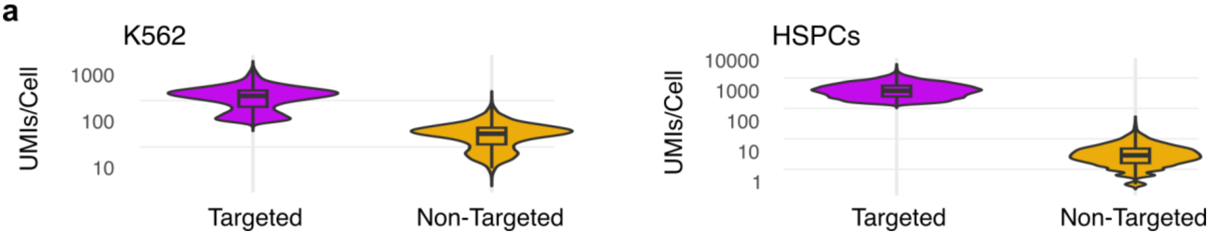


**Supplementary Fig. 5 | UMI distributions of mRNA target capture. a** Targeted gene capture in K562 cells and primary HSPCs shows strong enrichment of target genes relative to non-targeted transcripts.

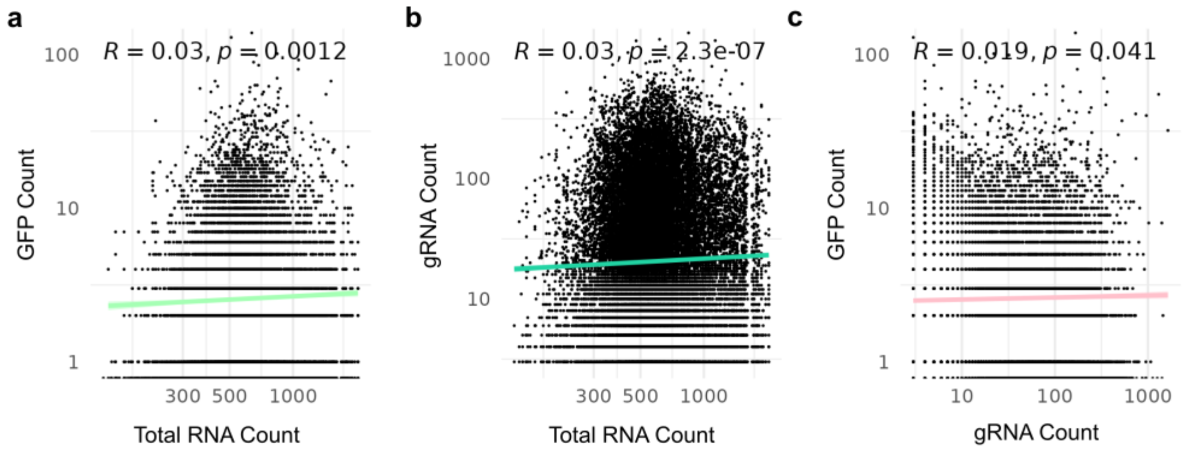


**Supplementary Fig. 6 | Correlations between total RNA UMI count, guide and GFP UMIs in PoC library measured in K562 cells.** GFP UMIs show no significant correlation with total RNA UMI counts, indicating that enhancer activity measurements are not driven by overall transcriptional output or technical capture efficiency. **b** gRNA UMI counts display a weak but significant correlation with total RNA counts, suggesting a modest dependence of gRNA capture on overall technical capture efficiency. **c** GFP and gRNA UMI counts show no significant correlation, indicating minimal cross-talk between the Pol II–driven and Pol III–driven expression cassettes.

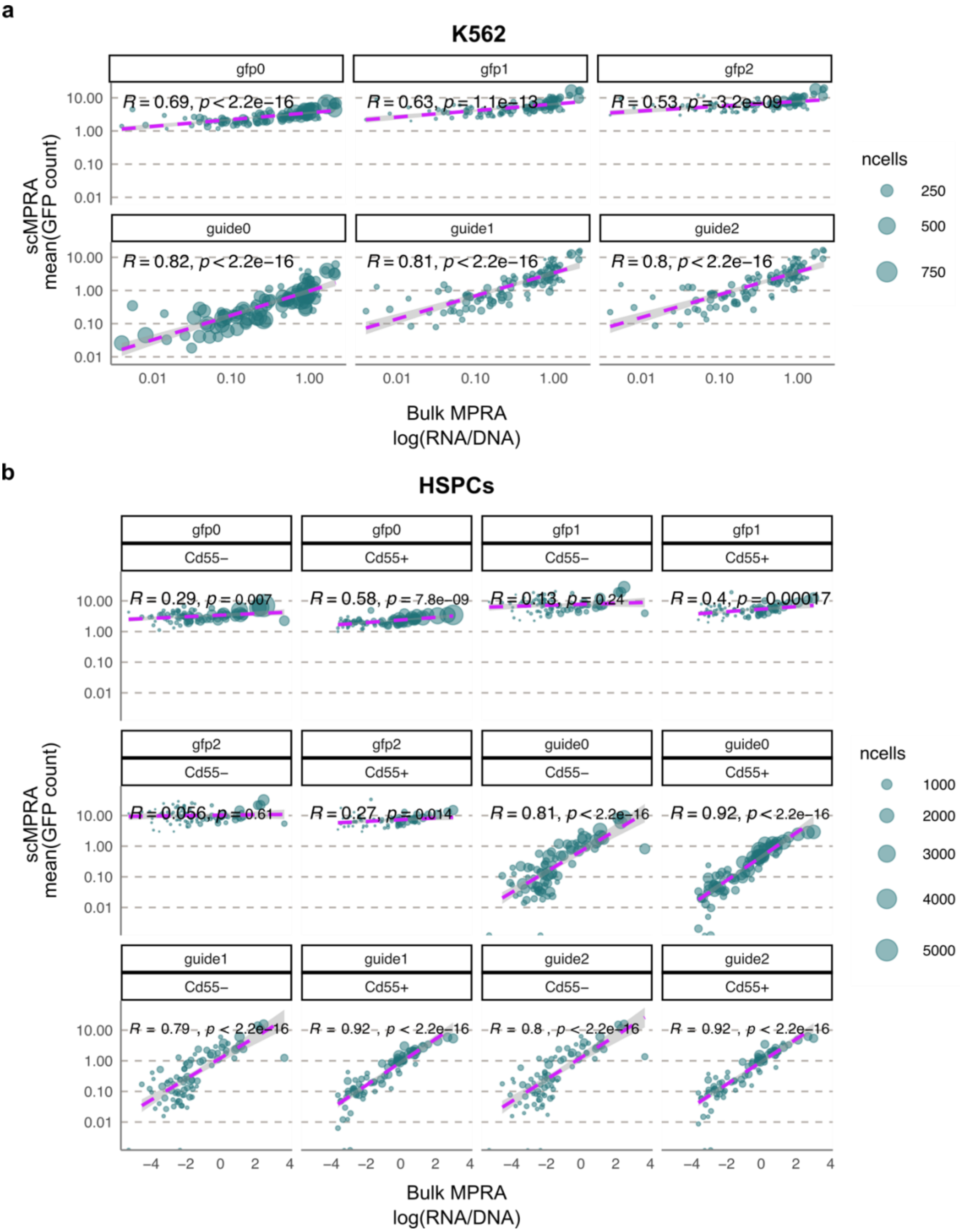


**Supplementary Fig. 7 | Comparisons of lenti-scMPRA preprocessing strategies and correlation with bulkMPRA data. a** $R^2$ values between enhancer activity estimates derived from lenti-scMPRA and bulk MPRA across different preprocessing strategies. GFP-only preprocessing quantifies enhancer activity directly from quantification (GFP) UMI counts, using a filter for minimum UMIs required of 0, 1 or 2 (gfp0, gfp1, gfp2). *Lenti-scMPRA-based* preprocessing first classifies cells as enhancer-positive based on presence-barcode (guide) detection thresholds (guide0, guide1, guide2 corresponding to presence barcode UMI > 0, >1, and >2, respectively), and subsequently quantifies enhancer activity using the corresponding GFP barcode counts. **b** Same analysis as in **a**, performed separately for two HSPC populations ($CD55^+$ and $CD55^-$).

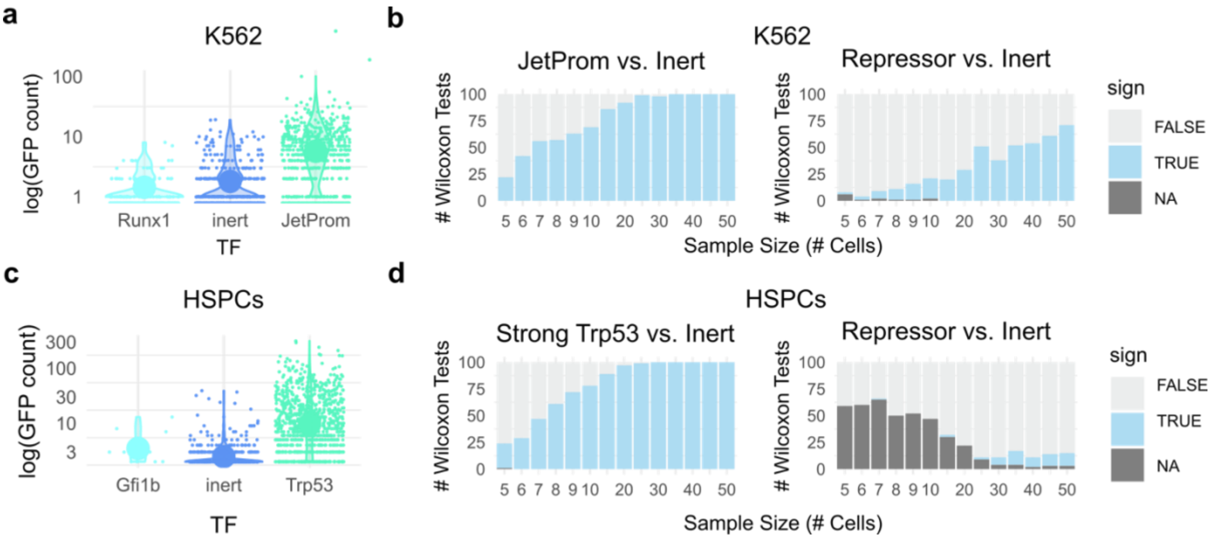


**Supplementary Fig. 8 | Sensitivity of sc-lentiMPRA. across K562 cells and HSPCs. a** Mean single-cell activity of inert random sequences, repressive Runx1 motif–containing sequences, and JeT promoter sequences in K562 cells. **b** Power analysis based on Wilcoxon rank tests in K562 cells showing the number of presence-barcode-positive single-cell measurements required to detect significant differences between JeT promoter activity sequences or repressive sequences and inert controls, respectively. Based on 100 pairwise Wilcoxon rank-sum tests with joint downsampling of all compared groups. **c** Mean single-cell activity of inert, repressive (Gfi1b-motif containing), and strong activator sequences (Trp53-motif containing) in HSPCs. **d** Power analysis in HSPCs, analogous to **b**, indicating the number of single-cell enhancer activity measurements required to detect significant activity differences between activator or repressive sequences and inert controls. NAs occur when both groups contain only zero values, resulting in no variance and preventing calculation of the Wilcoxon test statistic.

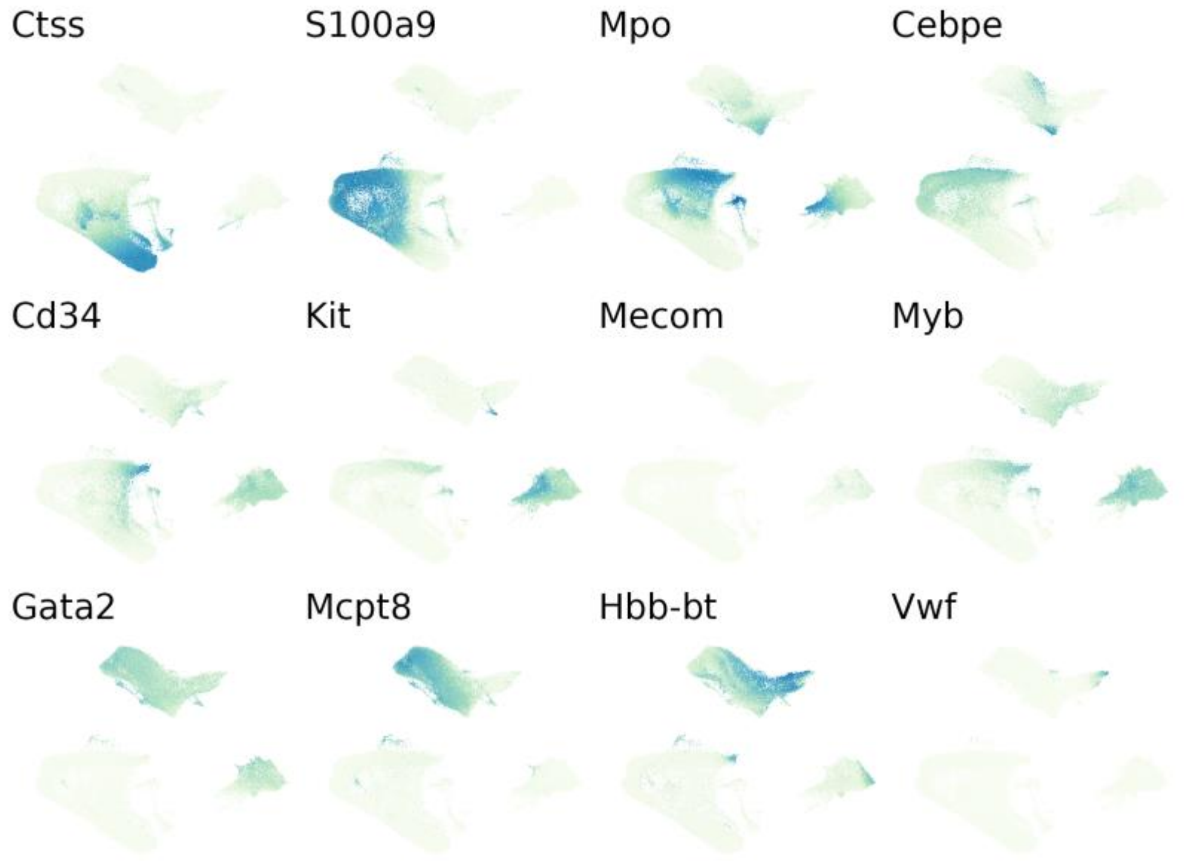


**Supplementary Fig. 9 | Marker gene expression of *in vitro* hematopoietic differentiation.** UMAP projection of the integrated single-cell dataset (Library Alpha, Library Beta) with cells colored by normalized expression of canonical markers spanning hematopoietic stem and progenitor states and downstream myeloid lineages. Shown are stem/progenitor markers **(**Cd34, Kit, Mecom, Myb, Gata2), granulocytic differentiation markers (Mpo, Cebpe), myeloid/monocytic markers (Ctss, S100a9), a mast cell/basophil-associated marker (Mcpt8), erythroid markers (Hbb-bt, Kel), and a megakaryocyte-associated marker (Vwf).

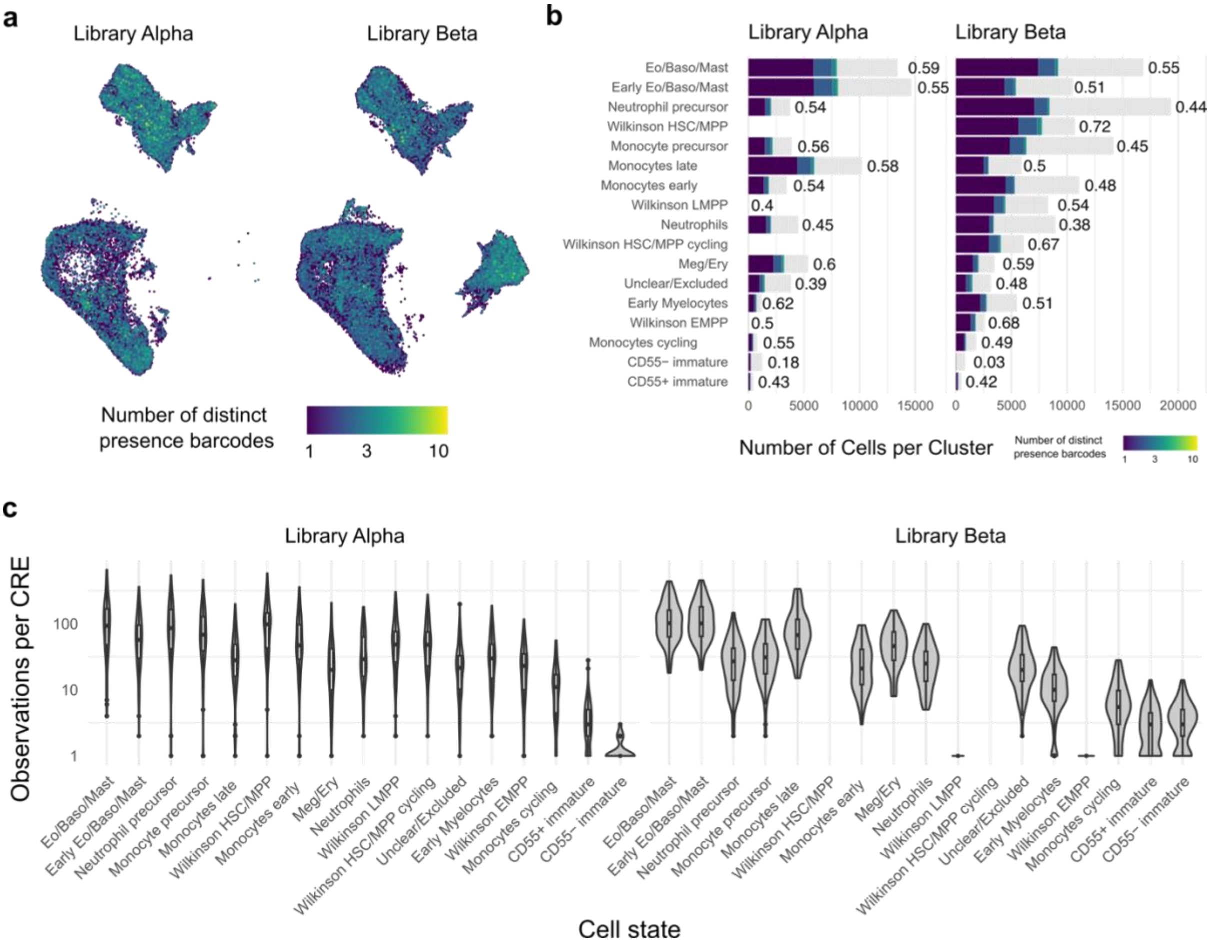


**Supplementary Fig. 10 | Efficacy of enhancer measurements across hematopoietic cell types. a** Low-dimensional integrated representation of all cells (Library Alpha, Library Beta) colored by the number of distinct presence barcodes detected using UMAP. **b** Proportions of cells containing at least one presence barcode across all cell types, indicating efficient and even construct delivery, as well as minimal silencing across lineages in both experiments. **c** Distribution of CRE coverage of the 84 candidate enhancers (Library Alpha), 80 candidate enhancers (Library Beta), respectively, per cell type.

**a**

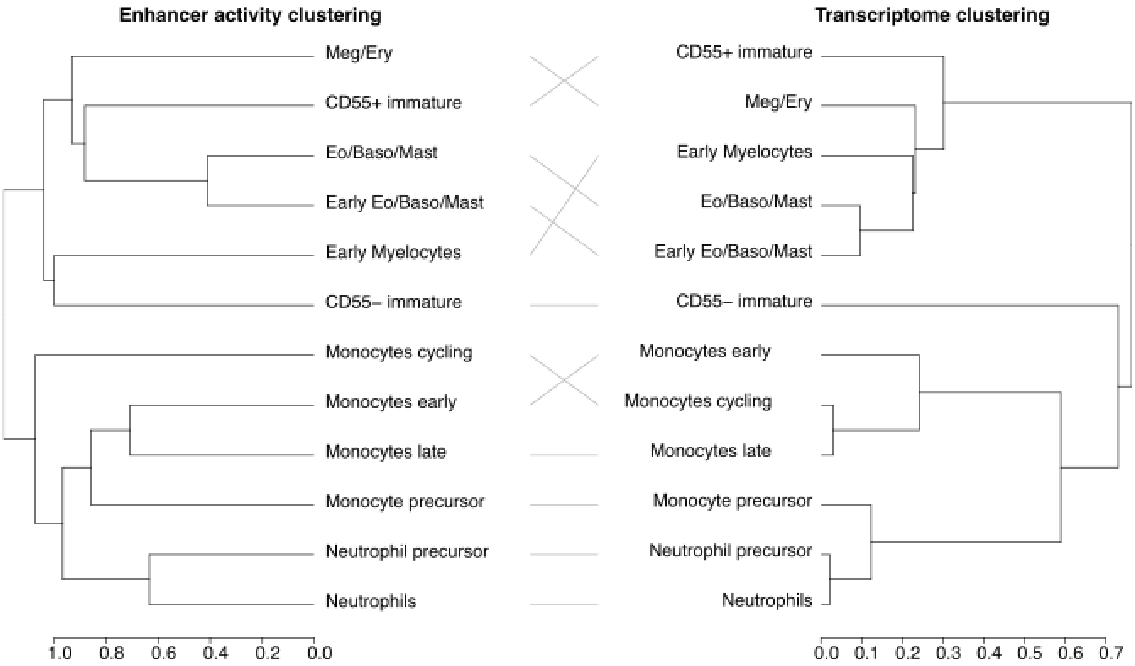


**Supplementary Fig. 11 | Transcriptome- and enhancer-based cell type clustering are highly concordant. a** Tanglegram comparing hierarchical clustering of cell types based on transcriptome profiles and enhancer activity profiles. The overall dendrogram structures indicate that enhancer activity patterns recapitulate clustering of transcriptome-based cell types.

**a**

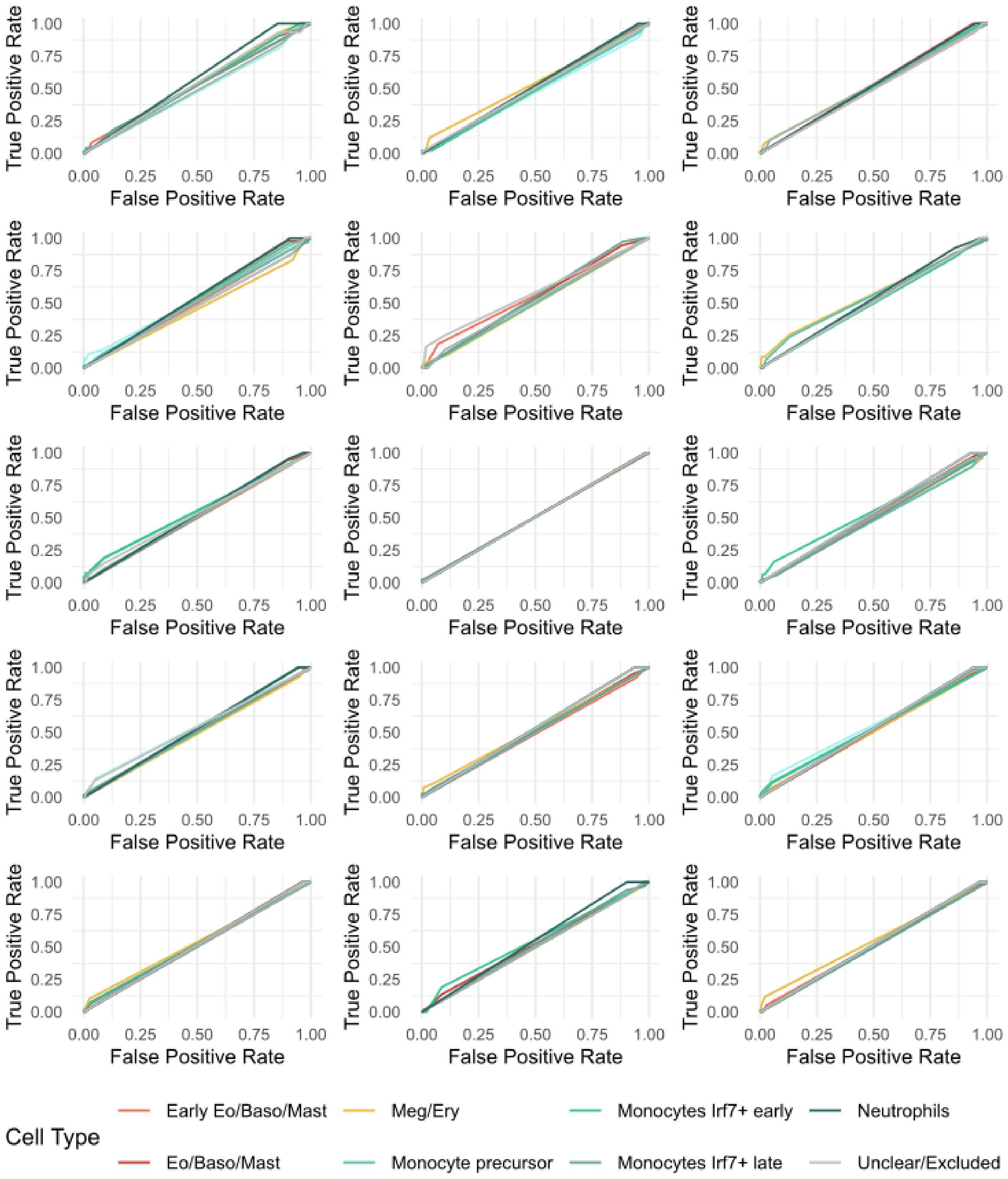


**Supplementary Fig. 12 | ROC curves of inert background control sequences. a** ROC curves of fifteen different inert background control sequences activate largely cell-type unspecific across different cell states.

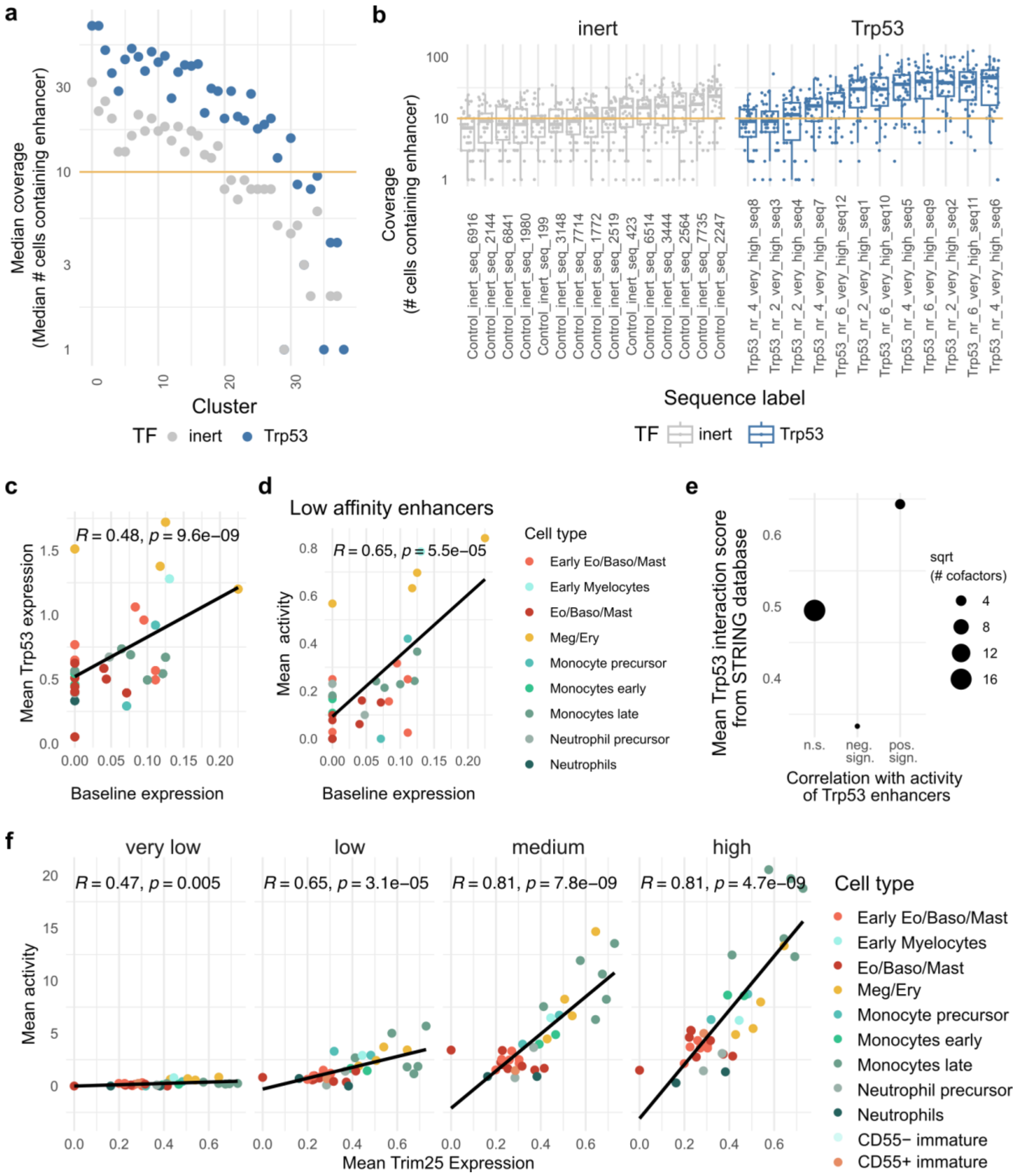


**Supplementary Fig. 13 | Trp53 enhancer-expression relationships. a** Median per-subcluster coverage for all Trp53 and inert control sequences (n = 38 subclusters obtained from clustering single-cell gene expression data). Y axis refers to the median number of cells per enhancer across all Trp53/control enhancers. **b** Number of observations (guide-positive cells) per Trp53 or inert control sequence across subclusters. **c** Correlation between mean Trp53 expression and baseline activity (inert control sequences), averaged per subcluster. **d** Correlation between baseline activity and mean activity of low-affinity Trp53 enhancers, indicating that the observed Trp53 enhancer-expression relationship is not explained by baseline activity differences. **e** STRING interaction scores between Trp53 and a curated set of transcriptional coregulators. Factors significantly correlated with Trp53enhancer activity exhibit higher mean interaction scores with Trp53 compared to non-significant or negatively correlated factors. **f** Correlation between Trim25 expression and Trp53 enhancer activity across affinity classes, highlighting Trim25 as the strongest positively associated cofactor candidate.

**a**

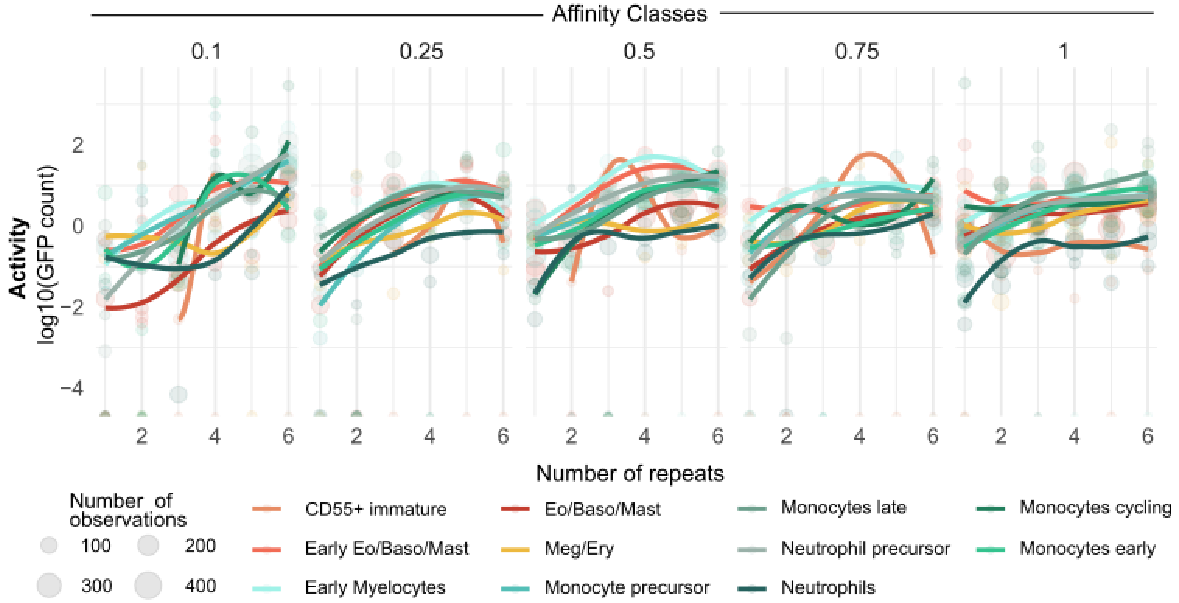


**Supplementary Fig. 14 | C/EBP enhancer activity across five affinity classes.** Enhancer activity of synthetic Cebpa enhancers across five affinity classes during early hematopoietic differentiation.

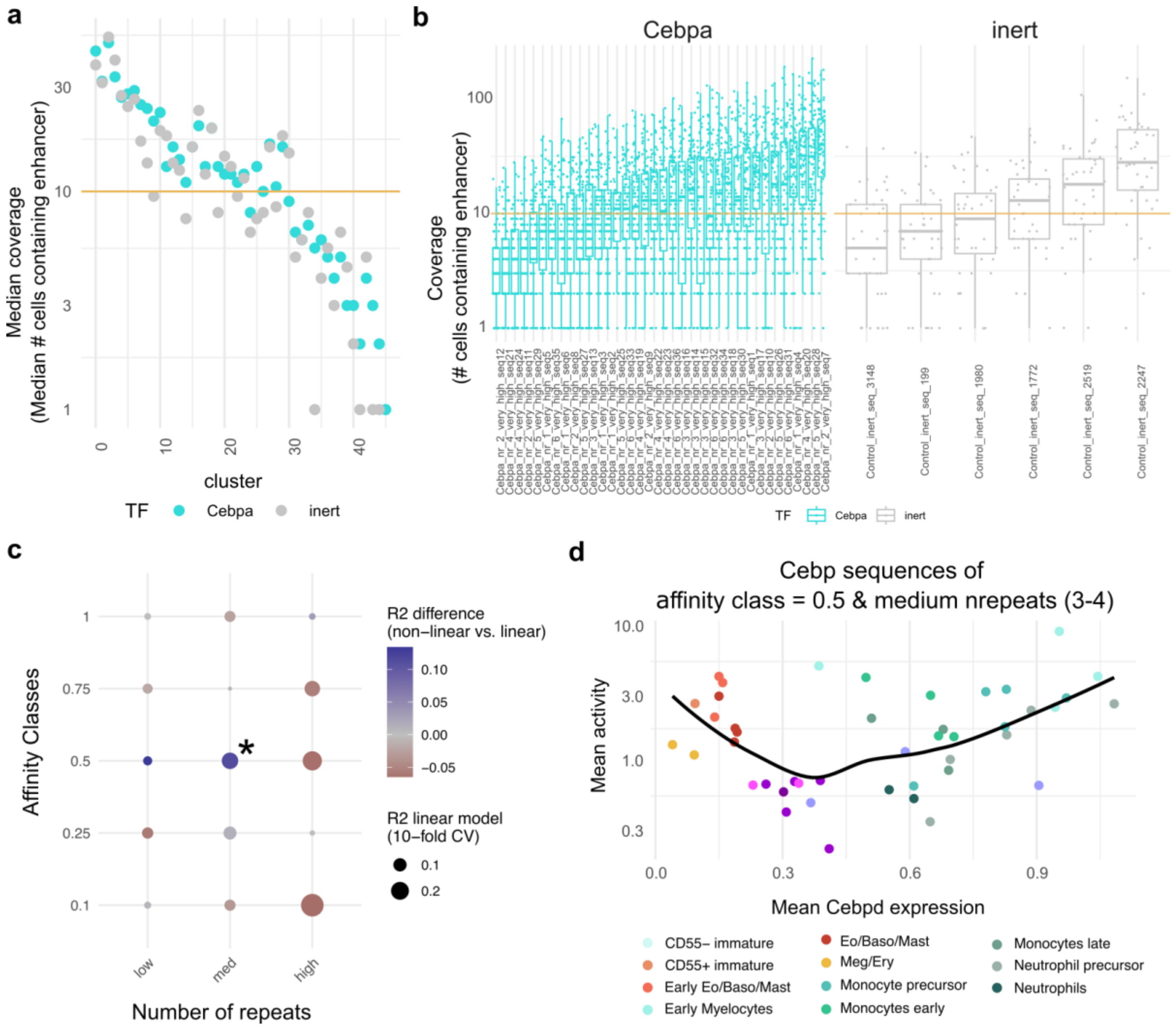


**Supplementary Fig. 15 | Cebp enhancer-expression relationships. a** Median per-subcluster coverage for all Cebp sequences and inert control sequences (n = 45 subclusters). **b** Number of observations (guide-positive cells) per Cebpa or inert control sequence across subclusters. **c,d** Evidence for a non-linear relationship between Cebp factor expression and enhancer activity. **c.** Enhancer activity was modeled with a linear model of Cebpa + Cebpad + Cebpe expression, and the coefficient of determination ($R^2$) was determined using 10-fold cross validation. Then, activity was modeled using project pursuit regression (PPR) of Cebpa + Cebpad + Cebpe expression, and $R^2$ was quantified using 10-fold cross validation. In PPR, a linear combination of explanatory variables is passed through a non-linear smoothing function. Color scale indicates mean difference in R2 between the linear and the PPR model across 10 independent 10-fold cross validation schemes. *: sequences shown in d. **d** Example of non-linear relationship between Cebpd expression and mean activity of Cebp enhancer sequences.
